# Lysosomal lipid metabolism promotes tumor cell invasion through local energetics and membrane lipid remodeling

**DOI:** 10.64898/2026.06.05.729624

**Authors:** Roseanne E. Nooren, Katherine M. Johnson, Maxwell G. Marley, Cody N. Rozeveld, Daniel F. Gibbard, Zinnarky K. Ortiz Correa, Mustafa Emre Gedik, Tianna Espe, Taro Hitosugi, Ian R. Lanza, Douglas G. Brownfield, Jun Liu, Mark A. McNiven, Gina L. Razidlo

## Abstract

Metabolic vulnerabilities in cancer have been targeted primarily to suppress tumor growth, but less is known about the metabolic requirements for tumor cell invasion. Here we report that lipid catabolism by cytosolic and lysosomal lipases supports pancreatic cancer cell invasion through both overlapping and distinct functional and metabolic mechanisms. Lysosomal acid lipase (LAL)-dependent lipid droplet catabolism promotes invadopodia formation and stabilization, enabling extracellular matrix degradation. In addition to modulating cellular energetics, lipidomics revealed that lipid droplet catabolism regulates cholesterol and membrane phospholipid levels. Using spatially resolved biosensors and cholesterol imaging, we found that lysosomal lipid catabolism occurs at invadopodia and sustains local ATP and membrane cholesterol. These findings identify spatially organized lipid catabolism as a mechanism that couples local energetics and membrane remodeling during the earliest steps of pancreatic cancer cell invasion.

## Introduction

Therapeutic strategies for cancer commonly target proliferating cancer cells. However, recent studies suggest the metabolic demands and dependencies of invasive cells may be distinct from proliferating cells (*1–5*). Thus, current strategies to target tumor growth may overlook invasive and metastasizing cells. At present, there is a critical need to understand the metabolic dependencies and drivers of invasive cell biology to target this dynamic and evasive cell population.

Pancreatic ductal adenocarcinoma (PDAC) is a deadly disease with a five-year relative survival rate of only 13% (*6*). Notably, pancreatic cancers are denoted by both early dissemination and distinctive metabolic rewiring (*7–10*). While the PDAC oncogenic driver KRAS has been strongly implicated in metabolic rewiring of glycolysis, glutamine dependence, and use of the pentose phosphate pathway, there is an emerging role for lipid metabolism in PDAC tumor growth and particularly metastasis (*11–19*). Our prior data suggested that excess lipids are stored in PDAC cells, and that upon migration tumor cells undergo a metabolic shift towards lipid catabolism and oxidative phosphorylation to fuel invasive migration (*14*).

Cells store excess lipids in the form of triglycerides and cholesterol esters in dynamic organelles called lipid droplets. Lipid droplets can be targeted by cytosolic lipases including adipose triglyceride lipase (ATGL) and hormone-sensitive lipase (HSL) to hydrolyze neutral lipids and liberate fatty acids (*14, 20*). In addition to cytosolic lipolysis, growing evidence reveals that lipid droplets can be catabolized by lysosomes via lipophagy. During lipophagy, lipid droplets are degraded by lysosomes by either autophagic engulfment or direct lysosomal contact (*21, 22*). The enzyme lysosomal acid lipase (LAL) liberates cholesterol and triglycerides from lipoproteins and hydrolyzes cholesterol esters and neutral lipids from lipid droplets (*22–25*). The relative metabolic and functional contributions of cytosolic versus lysosomal lipid catabolism are poorly understood. As lysosomal activity has been shown to be elevated in pancreatic cancer cells to support proliferation, we hypothesized that lipophagy is also active in pancreatic cancer cells to support lipid droplet catabolism.

Here, we show that lysosomal and cytosolic lipases contribute to lipid droplet dynamics and promote tumor cell invasion in pancreatic cancer cells through distinct mechanisms. Whereas the cytosolic lipase HSL promotes cell migration, LAL is required for pancreatic cancer cell invasion and invadopodia-mediated degradation of the extracellular matrix. Further, live cell imaging and a lipid droplet mistrafficking strategy reveal the importance of localized lipid droplet catabolism within PDAC cells. Mechanistically, lipid droplet catabolism fuels tumor cell invasion by regulating mitochondrial cellular energetics and membrane lipid dynamics, and LAL activity contributes to levels of ATP and membrane cholesterol specifically at invadopodia. These findings help to demonstrate the importance of localized lipid droplet catabolism via lysosomal acid lipase in cancer cell invasion.

## Results

### Lipid droplet catabolism through lysosomal acid lipase in PDAC cells

Lipolysis of stored lipid droplets has been shown to support pancreatic cancer cell invasion, though prior research has focused on cytosolic lipases (*14*). As lysosomal biogenesis and activity are amplified in pancreatic cancers, we sought to determine the contributions of lysosomal catabolism of lipid droplets in pancreatic cancer cells (*26–28*). We first evaluated the expression levels of LAL, gene name *LIPA,* at the mRNA level in pancreatic adenocarcinoma tumors. Interestingly, there was a significant increase in *LIPA* expression in pancreatic adenocarcinomas from the TCGA database compared to GTEx normal pancreas (Figure 1A) (*29, 30*). We also noticed a significant increase in *LIPA* mRNA expression in tumor samples compared to matched normal pancreas (GSE 15471, Figure 1B) (*31*). We hypothesized that amplified *LIPA* expression in pancreatic cancers indicates a role for lipid droplet catabolism by lysosomes. This led us to further investigate the role of LAL in pancreatic cancer cells.

**Figure 1.**
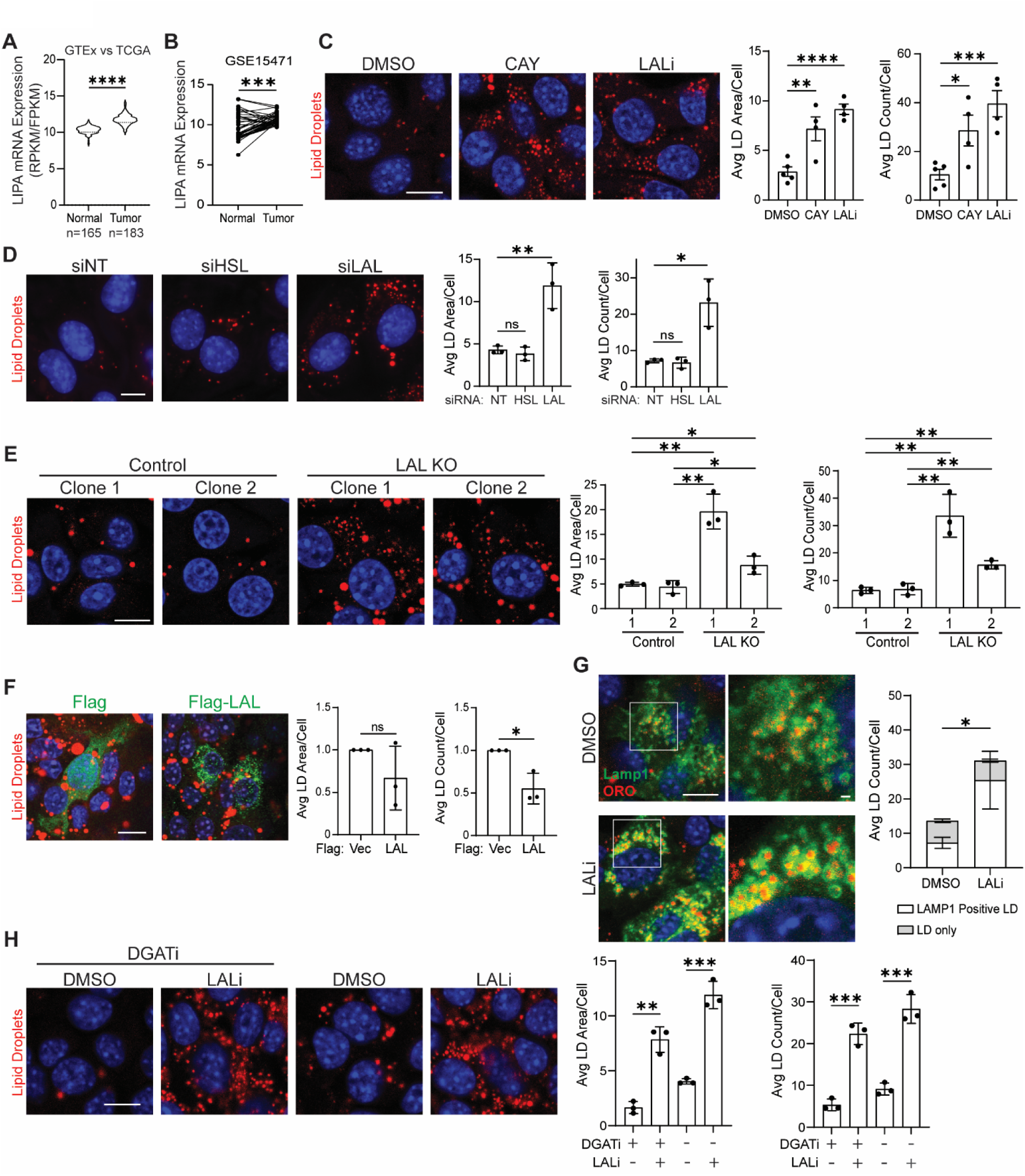
Lysosomal acid lipase catabolizes lipid droplets in PDAC cells. **A)** Relative *LIPA* mRNA expression in normal healthy pancreas (GTEx, 165 cases) compared to pancreatic cancer tumors (TCGA-PAAD, 183 cases). **B)** *LIPA* mRNA expression profile from GSE15471 data set of paired 39 PDAC tumor and normal adjacent pancreas samples. p value was calculated using paired Student’s t test. **C)** mKPC cells treated with DMSO (vehicle control), CAY10499 (CAY, 10μM), or LAListat1 (LALi, 50μM) for 24h were stained for lipid droplets (Oil Red O, red) and nuclei (Hoechst, blue). **D)** mKPC cells transfected with control non-targeting (NT) siRNA, HSL siRNA, or LAL siRNA and stained for lipid droplets (red) and nuclei (blue). **E)** Staining of lipid droplets (red) and nuclei (blue) of LAL Knockout (KO) mKPC cells compared to control cells. **F)** mKPC cells transfected with Flag-LAL or Flag empty vector (control), stained for Flag (green), lipid droplets (red) and nuclei (blue). 20 fields were imaged per experiment per condition for each biological replicate. **G)** mKPC cells transfected with LAMP1-GFP were treated with DMSO or 50μM LALi for 24h and stained for lipid droplets (red). Lipid droplets were scored manually as being LAMP1 positive or negative. Statistical significance was calculated for LAMP1-positive lipid droplets. **H)** mKPC cells were treated with DGATi 1 and 2 (10µM) with DMSO or 50μM LALi for 24h. **(C, D, E, F, and H)** Representative images with quantitation of average lipid droplet area per cell and average number of lipid droplets per cell from at least three independent biological replicates. Unless otherwise indicated, 10 fields were imaged per experiment for each biological replicate. Scale bar = 10µm; insets (G) magnified images scale bar = 1µm. p values were calculated using unpaired Student’s t test: *p < 0.05, **p < 0.01, ***p < 0.001 ****p < 0.0001, ns = not statistically significant.

We first examined the requirement for LAL in comparison with cytosolic lipases in lipid droplet turnover in pancreatic cancer cells. LAListat1 (LALi) is a chemical inhibitor of LAL, and CAY10499 (CAY) reduces cytosolic lipase activity of HSL and monoacylglycerol lipase (MGL). To visualize lipid droplet content, three human PDAC cell lines (DAN-G, PANC-1 and BxPC-3) and one mouse PDAC cell line (mKPC) were stained with Oil Red O. Inhibition of either LAL or HSL/MGL led to a marked increase in average lipid droplet number and average lipid droplet area per cell, implicating both lysosomal and cytosolic lipolysis in lipid droplet breakdown (Figure 1C, Figure S1A-C). RNAi-mediated knockdown of HSL or LAL similarly increased the average lipid droplet area and number per cell in the mKPC and DAN-G cells (Figure 1D, Figure S1D-H). We then generated LAL knockout mKPC cell lines using CRISPR-Cas9. In two different clones with genetically distinct profiles, we observed a marked increase in lipid droplet content compared to the control cells, indicative of disrupted lipid droplet catabolism (Figure 1E). Furthermore, overexpression of LAL in mKPC led to a dramatic reduction in average lipid droplet number per cell, congruent with amplified lipid droplet catabolism (Figure 1F). As demonstrated previously in our laboratory, overexpression of HSL in mKPC and iKras cells similarly lead to a decrease in lipid droplet content per cell due to enhanced catabolism (*14*). Taken together, these approaches demonstrate that LAL and cytosolic HSL/MGL mediate lipid droplet catabolism all contribute to overall lipid droplet levels in PDAC cell lines.

To indicate lipid droplet catabolism more specifically in lysosomes upon LAListat1 treatment, we visualized lipid droplets with Oil Red O staining and lysosomes by overexpressing the lysosomal marker LAMP1-GFP. Chemical inhibition of LAL increased the percentage of lipid droplets within lysosomes by 2-3 fold compared to DMSO control treatment (Figure 1G, Figure S1I). To ensure that lipid droplet accumulation specifically indicated impaired lipid droplet catabolism instead of promoting triglyceride synthesis, we blocked synthesis of new lipid droplets by inhibiting diacylglycerol transferases (DGAT) 1 and 2 by using two DGAT inhibitors (DGATi). DGAT 1 and 2 convert diacylglyceride into triglyceride in the last step of triglyceride synthesis for storage in lipid droplets. In the presence of DGATi, while overall lipid droplet content was reduced, LALi still induced a significant increase in the mean lipid droplet number and mean lipid droplet area in mKPC and DAN-G cells (Figure 1H, Figure S1J). This suggests lipid droplet accumulation following inhibition of LAL or HSL/MGL is not due to increased lipogenesis, but rather impaired lipid droplet catabolism.

### Lysosomal lipolysis promotes cancer cell invasion

We next investigated the requirements for lysosomal and cytosolic lipolysis in the support of several key cellular processes. Cell viability was measured utilizing a CCK8 assay, but inhibition of neither LAL nor HSL/MGL reduced cell viability under steady state conditions in culture (Figure S2A). We have previously shown the action of cytosolic lipases including HSL promote tumor cell invasion (*14*). Therefore, we performed multiple functional assays to examine the contributions of cytosolic versus lysosomal lipolysis to migration and invasion of pancreatic cancer cells. Transwell assays revealed that both cytosolic and lysosomal lipolysis promotes cancer cell invasion. Pharmacological inhibition of HSL/MGL using CAY10499 or LAL using LAListat1 markedly reduced transwell invasion of multiple PDAC cell lines by up to 90% (Fig 2A-B, Figure S2B). Similarly, RNAi-mediated knockdown of HSL or LAL in mKPC and DAN-G cells also reduced tumor cell invasion (Figure 2C-H). Conversely, overexpression of LAL enhanced transwell invasion in mKPC cells by 50% (Figure 2I). These data support a role for both cytosolic and lysosomal lipases in promoting tumor cell invasion, though the relative contributions of each lipase varied by cell line.

**Figure 2.**
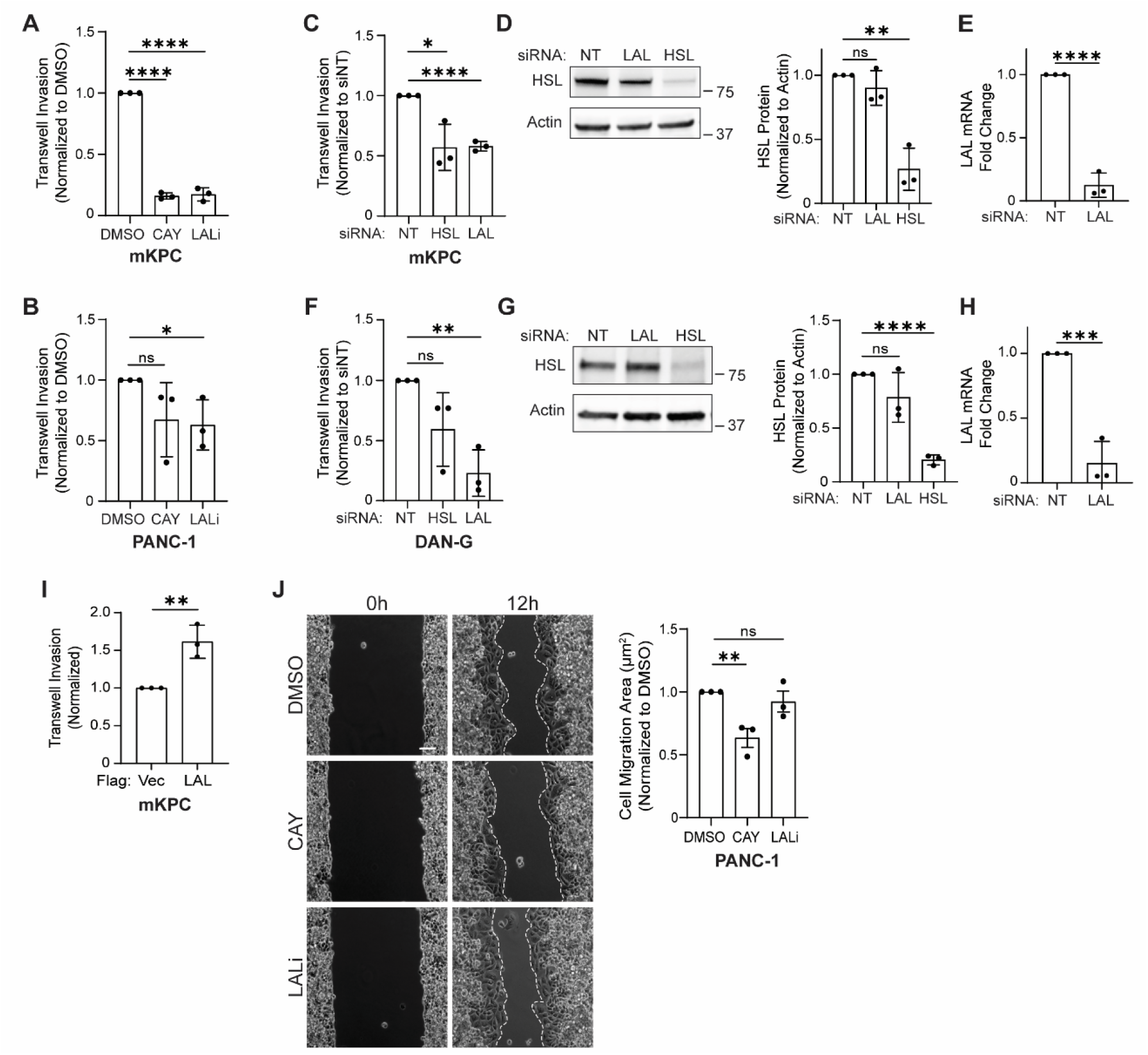
Lysosomal lipase activity promotes tumor cell invasion. **A-B)** Quantitation of transwell cell invasion by mKPC **(A)** and PANC-1 **(B)** cells after chemical inhibition of HSL/MGL (CAY, 10μM) or LAL (LALi, 50μM) for 24h (mKPC) or 48h (PANC-1). **C)** mKPC cells transfected with control non-targeting (NT), HSL, or LAL siRNAs, and were seeded in a transwell assay for 24h. **D)** Representative immunoblot and quantitation for three independent replicates show the degree of HSL knockdown. **E)** LAL mRNA levels were measured by qPCR from three independent replicates to confirm LAL knockdowns. **F)** DAN-G cells transfected with control non-targeting (NT), HSL, or LAL siRNAs, and were seeded in a transwell assay for 24h. **G)** Representative immunoblot and quantitation for three independent replicates show the degree of HSL knockdown. **H)** LAL mRNA levels were measured by qPCR from three independent replicates to confirm LAL knockdowns. **I)** mKPC cells transfected with Flag-LAL or Flag empty vector control and seeded in a transwell assay for 24h. Flag epitope staining confirmed that only Flag expressing cells were quantified for three independent biological replicates. **J)** Representative images of wound healing cell migration at 0h and 12h in PANC-1 cells pre-treated with DMSO, 10μM CAY, or 50μM LALi for 24h. Quantitation of cell migration area (μm^2^) for 3 independent biological replicates (3 fields per timepoint for each condition). Scale bar = 10µm. Throughout, graphed data represent the mean ± SD for at least three independent biological replicates. p values were calculated using unpaired Student’s t test. *p < 0.05, **p < 0.01, ***p < 0.001 ****p < 0.0001, ns = not statistically significant.

We subsequently tested how LAL and HSL/MGL impacted cancer cell invasion within the context of lipid overloading. After pretreating cells with oleic acid, inhibition of either HSL/MGL or LAL under washout conditions substantially increased lipid droplet retention by two-fold (Figure S2C-D). Surprisingly, while chemical inhibition of HSL/MGL still reduced PDAC cell invasion in oleate-loaded cells, chemical inhibition of LAL had no effect on cancer cell invasion following oleate loading (Figure S2E). Since there was a substantial decrease in invasion under basal lipid conditions, this suggests that it is not solely the accumulation of lipid droplets or catabolism that promote invasion. Rather, this suggests that specific LAL-dependent fatty acid species may modulate invasive biology. Furthermore, this indicated functional differences between the roles of cytosolic lipolysis and lysosomal lipid catabolism in cancer cell invasion.

### Lysosomal lipolysis regulates invadopodia formation to promote ECM degradation

Transwell invasion assays recapitulate multiple steps required for invasive dissemination *in vivo*, including chemotaxis, directional migration, and pro-invasive degradation of the extracellular matrix. To dissect the process of invasion and define the specific contributions of lipid droplet catabolism, we next examined steps of the invasive process. Two-dimensional wound healing assays were used to evaluate cell migration, which requires migratory polarization, actin dynamics, and membrane protrusions, but does not require degradation or remodeling of the extracellular matrix. As mentioned above, inhibition of cytosolic HSL using CAY10499 reduced two-dimensional cell migration (Figure 2J, Figure S2F). Surprisingly, inhibition of lysosomal acid lipase through LAListat1 had no effect on two-dimensional cell migration, although it potently impacted transwell invasion (Figure 2J, Figure S2F).

A critical difference between 2D cell migration and invasion is the presence of extracellular matrix (ECM). Cancer cells degrade and remodel ECM to disseminate away from the primary tumor. To investigate if lysosomal acid lipase is important for degradation of the ECM *in vitro*, we plated DAN-G cells on a fluorescently conjugated gelatin matrix (*32*). Whereas DAN-G cells exhibited a robust degradation of ECM as quantified by loss of the fluorescent matrix, inhibition of lysosomal acid lipase significantly decreased the area of ECM degradation by 50%. While the HSL inhibitor CAY10499 also reduced matrix degradation, it was to a lesser degree than LAL inhibition (Figure 3A-B, Figure S3A-B).

**Figure 3.**
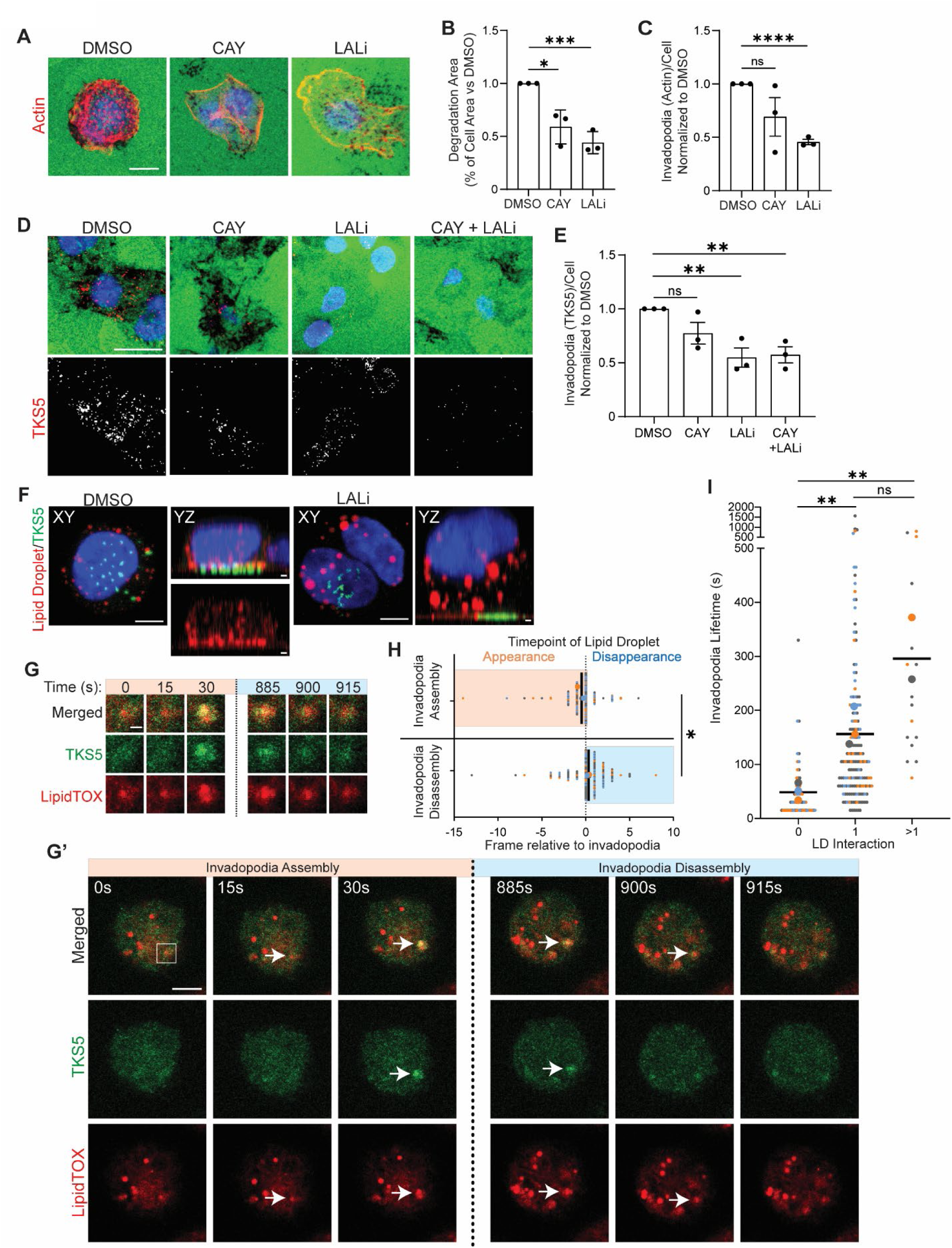
Lysosomal acid lipase activity promotes invadopodia-mediated extracellular matrix degradation. **A)** DAN-G cells treated with DMSO, 10μM CAY, or 50μM LALi for 24h were seeded on Oregon Green gelatin-coated coverslips and stained for actin (Phalloidin, red) and nuclei (Hoechst, blue). At least 110 cells were scored per condition in each of three biological replicates. Scale bar = 10µm. **B)** Quantitation of average gelatin degradation area normalized to cell area. **C)** Quantitation of puncta positive for both actin and gelatin degradation per cell. **D)** DAN-G cells treated as in (A) were stained for endogenous TKS5 to detect invadopodia (red) and nuclei (Hoechst, blue). Scale bar = 20µm. **E)** Quantitation of average TKS5-positive invadopodia number per cell after inhibitor treatment. At least 240 cells were scored per condition in each of three independent biological replicates. **F)** Z stack orthogonal projections on XY plane and YZ plane of DAN-G cells transfected with TKS5-GFP (green) and stained for lipid droplets (Oil Red O, red) and nuclei (Hoechst, blue) treated with DMSO or 50μM LALi for 24h. Scale bar XY = 5µm; Scale bar YZ = 1µm. **G-I)** Invadopodia lifetime for DAN-G cells transfected with TKS5-GFP (invadopodia, green) and stained with LipidTOX Deep Red (LD, red) and plated on unlabeled gelatin for 5h. **(G’)** Representative images shown were from the indicated timepoints. Arrows point to TKS5 and lipid droplet presence. The invadopodia and lipid droplets were maintained in the frames not shown between 30 and 885s. **(G)** Magnified inset shows the boxed region where lipid droplets and invadopodia are colocalized. Larger image scale bar = 5µm; magnified image scale bar = 1µm. **(H)** Graph of LD presence relative to invadopodia. The timepoint of Invadopodia appearance (orange) and disappearance (blue) is set to t=0 and lipid droplet detection was measured relative to this point. p values were calculated using paired Student’s t test **(I)** TKS5-GFP positive invadopodia lifetime was quantitated and graphed based on number of lipid droplet interactions. Data points indicate all individual invadopodia imaged over three independent experiments. Graphed data represent the mean ± SD for at least three independent biological replicates unless otherwise indicated. p values were calculated using unpaired Student’s t test unless otherwise indicated. *p < 0.05, **p < 0.01, ***p < 0.001 ****p < 0.0001, ns = not statistically significant.

Invadopodia are F-actin based protrusions that target sites for proteinase secretion, leading to degradation of the extracellular matrix and invasion into the surrounding tissue (*33, 34*). Exogenous lipids have been linked to invadopodia formation in breast cancer cells (*35*). Consequently, we evaluated if cytosolic or lysosomal lipases support invadopodia formation or function by staining for actin or the invadopodial scaffolding protein TKS5. LAListat1 treatment significantly decreased the number of actin puncta colocalized with ECM degradation sites by 50% (Figure 3C) as well as invadopodial TKS5 puncta (Figure 3D-E). In contrast, HSL/MGL inhibition did not significantly reduce the number of invadopodia. This demonstrates that lysosomal lipid catabolism supports the formation or function of invadopodia in the PDAC cells to actively degrade the extracellular matrix.

Interestingly, confocal microscopy and Z stack reconstruction suggested the colocalization of lipid droplets with some invadopodia at the base of the cell (Figure 3F). However, this interaction appeared disrupted in the LAListat1-treated cells, which have fewer invadopodia. To determine if lipid droplet localization to invadopodia impacts invadopodia formation or dynamics, we performed live cell imaging in cells expressing TKS5-GFP to label invadopodia and LipidTOX Deep Red to label lipid droplets. Quantitation of frame-by-frame interactions between lipid droplets and invadopodia revealed lipid droplets present before the formation of TKS5 positive puncta, during invadopodia lifetime, and after dissolution (Figure 3G-H, Supplemental Movie 1-2). In most cases of colocalization, the lipid droplet appeared first prior to invadopodia formation, and the lipid droplet disappeared from view after invadopodia disassembly (Figure 3H). Notably, colocalization with a lipid droplet significantly increased invadopodial lifetime, including interaction with one lipid droplet or multiple lipid droplets (Figure 3I). TKS5-positive puncta negative for lipid droplet interactions were highly dynamic and had an average lifetime of only 46 seconds. In contrast, interactions with one lipid droplet increased the mean invadopodial lifetime to over 169 seconds in the DMSO condition, suggesting contributions of lipid droplets to invadopodia stability. In LAListat1-treated cells, the overall number of invadopodia was decreased and the number of lipid droplets were substantially increased, as shown above (Figure 3A-E). Of the TKS5 invadopodia that formed in the LAListat1 treated cells, association with a lipid droplet similarly increased invadopodia lifetime from an average of 111 seconds without lipid droplet interactions, to 289 seconds with lipid droplet interactions (Figure S3C-D). Of the fewer invadopodia that were detectable under LAListat1 treatment, there was still increased invadopodia stability if adjacent to a lipid droplet. Together, these data suggest that invadopodia that form near lipid droplets have a longer lifetime and indicate that catabolism of lipid droplets is important in invadopodia formation and stabilization.

### Spatial regulation of lysosomal acid lipases promotes invasion

Lysosomes have previously been shown to have a regulatory role in invadopodial biology and invasive matrix degradation (*36*). The requirement for LAL activity in invadopodia number and the intriguing localization of invadopodia near lipid droplets led us to speculate that lipophagy could be occurring at or near invadopodial sites. To test this, we expressed a fluorescent biosensor for lipophagy, GFP-RFP-PLIN2 (Figure 4A) (*21, 37*). The lipid droplet protein PLIN2 (Perilipin-2) is fused to both GFP and RFP, with both red and green signals detected when the lipid droplets are in the cytosol. If any or all of the lipid droplet is internalized into the lysosome for lipophagy, the acidic environment of the lysosome quenches the GFP fluorescence. This leaves the lipid droplet labelled by only RFP. Strikingly, we observed at a single fixed timepoint that 15% of TKS5-positive invadopodia were also positive for sites of lipophagy (Figure 4A-B). This led us to hypothesize that lipid droplet catabolism might be spatially regulated to promote pancreatic cancer cell invasion.

**Figure 4.**
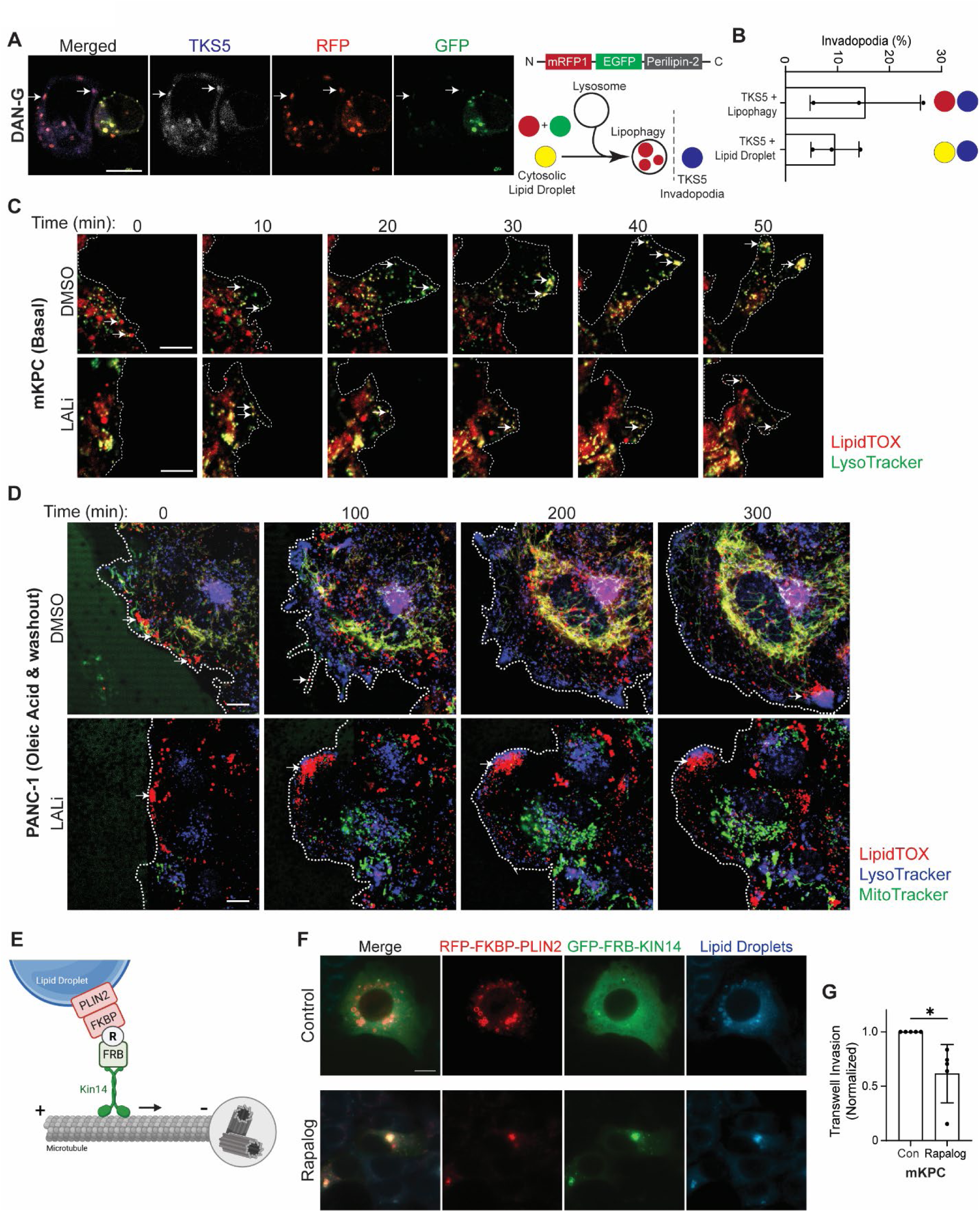
Lipid droplet localization is critical for tumor cell invasion. **A)** DAN-G cells transfected with RFP-eGFP-PLIN2 and stained for TKS5 (blue, pseudo colored to gray). Arrows indicate TKS5 + RFP puncta. Scale bar = 10µm**. B)** Quantitation of average number per cell of blue and red only puncta (lipid droplets within lysosomes at TKS5 invadopodia), and blue, red and green puncta (cytosolic lipid droplets at TKS5 invadopodia). Representative images and quantitation for three independent biological replicates; n=20 fields for each replicate. **C)** Representative images of mKPC cells treated with DMSO or 50μM LALi for 24h and stained with LysoTracker (lysosome, green) and LipidTOX (lipid droplet, red) showing lysosome and LD localization at the periphery of the cell during wound healing migration. Live cell images taken at the indicated timepoints. Scale bar = 10µm. **D)** Representative images of PANC-1 cells treated with oleic acid (200µM), then washed out and fresh media added with either DMSO or 50μM LALi and stained with LysoTracker (blue), LipidTOX (red) and MitoTracker (green). Lipid droplet (red) localization shown at the periphery of the cell during wound healing migration. Live cell images taken at the indicated timepoints. **E)** Schematic of lipid droplet redistribution system. The PLIN2-FKBP fusion protein (red) labels lipid droplets (blue). Addition of the Rapalog A/C heterodimerizer (R) tethers lipid droplets to the FRB-KIN14 fusion protein (green) for retrograde trafficking. **F)** Representative images of mKPC cells transfected with RFP-FKBP-PLIN2 and GFP-FRB-KIN14, treated with or without Rapalog (10nM, 4h). Lipid droplets were stained with MDH (blue). **G)** mKPC cells treated as in **(F)** were seeded in a transwell assay for 24h. Only transfected cells indicating lipid droplet trafficking were scored. Scale bar = 10µm. Graphed data represents the mean ± SD for five independent biological replicates. p values were calculated using unpaired Student’s t test. *p < 0.05.

Separately, we observed examples of lipid droplets at the leading edge of cells migrating *in vitro* (Figure 4C-D, Figure S4A-B, Supplemental Movie 3-4). Particularly in the presence of LAListat1 to prevent lipid droplet degradation, lipid droplets could be detected at the cell periphery during wound healing assays (Figure 4C-D, Figure S4A). This was amplified even further upon loading with oleic acid to increase lipid droplet content (Figure 4D, Supplemental Movie 5-6). Live cell imaging indicated trafficking of lipid droplets in the direction of growing protrusions (Figure 4C-D). Of note, since lipid droplet catabolism and turnover occur during cell invasion (*14*), this observation was sensitive to the timepoint of the experiment and was difficult to capture, particularly at later stages.

The intriguing localization of lipid droplets at invadopodia and in the leading edge led us to hypothesize that lipid droplet localization to the invasive periphery supports tumor cell invasion. To test this, we employed a chemical dimerization system to control the trafficking of lipid droplets and mistraffic them away from peripheral protrusions (*38*). The lipid droplet protein PLIN2 was fused to FKBP (FK506-binding protein) and was co-expressed with a construct encoding FRB fused to the retrograde kinesin Kin14. The addition of the Rapalog A/C Heterodimerizer (AP21967) induces heterodimerization between FKBP and FRB, thus tethering the lipid droplets to the retrograde kinesin and trafficking them away from the cell periphery, resulting in perinuclear clustering (Figure 4E-F, Figure S4C)(*38*). Notably, mistrafficking of the lipid droplets led to a marked reduction in transwell invasion by nearly 50% (Figure 4G, Figure S4D). This intriguing finding suggests that the localization of lipid droplets, and potentially localized lipid droplet catabolism, contributes to tumor cell invasion. Notably, when the PDAC cells were pre-loaded with oleic acid and then washed out, lipid droplet mistrafficking did not inhibit invasion (Figure S4E), similar to the effects of LAListat1 and again suggesting it is not simply the presence of lipid droplets, but potentially indicating functional specificity to lipid content and metabolic output. Treatment with the Rapalog A/C Heterodimerizer itself in the absence of the engineered constructs had no effect on transwell invasion (Figure S4F). Together, these data indicate that not only is lipid droplet catabolism important, but lipid droplet localization at or near the sites of invasive protrusions is critical for tumor cell invasion.

### Lysosomal acid lipases promote ATP production and oxidative phosphorylation in PDAC

We next sought to determine how lipid droplet catabolism promotes invasion. We first investigated how lysosomal acid lipase impacts fatty acid oxidation, mitochondrial respiration, and cellular energetics. Fatty acid oxidation was measured by tracing ^13^C palmitic acid into tricarboxylic acid cycle metabolites. When comparing treatment with LAListat1 to DMSO in DAN-G cells, there was no significant difference in M+2 labelling of citrate (Figure S5A). However, this assay relies on incorporation of exogenous palmitic acid, which may not capture activity of lysosomal acid lipases in stored lipid droplets but indicates no global defect in fatty acid oxidation. Thus, we further examined the relative contributions of cytosolic and lysosomal lipases to mitochondrial oxidative metabolism using a Seahorse MitoStress assay (Figure 5A-D). Upon chemical inhibition of either LAL or HSL/MGL in DAN-G cells, we observed a similar and significant decrease in oxygen consumption rate for both basal and maximal respiration normalized to DMSO treatment (Figure 5A-C, Figure S5B-E). When both HSL/MGL and LAL were inhibited together, there was an additive inhibition of respiration in the DAN-G cells. This suggests that lysosomal acid lipase and cytosolic lipases can impact mitochondrial respiration potentially through parallel pathways.

**Figure 5.**
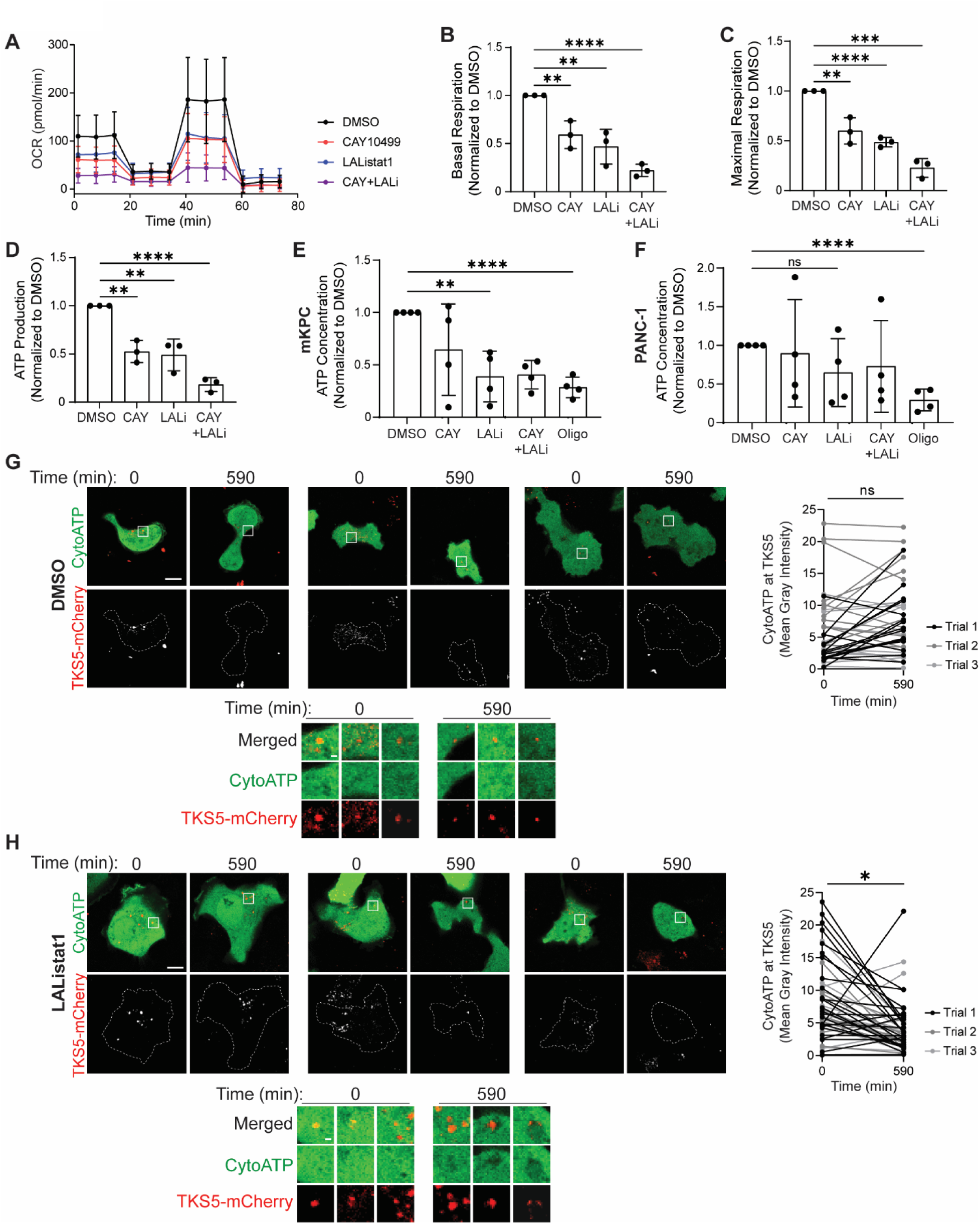
Cytosolic and lysosomal acid lipase activity regulates oxidative metabolism and ATP production. **A)** Quantitation of average normalized oxygen consumption rate (OCR) from Seahorse metabolic analysis for DAN-G cells treated with DMSO, 10μM CAY, 50μM LALi, or CAY+LALi for 24h. Data extracted from **(A)** includes basal respiration **(B)**, maximal respiration **(C)**, and ATP production **(D)**. Graphed data represents the mean ± SD for three independent biological replicates. **E-F)** Quantitation of ATP concentration using ATP Detection Luminescence assay kit for mKPC **(E)** and PANC-1 **(F)** cells treated with DMSO, 10μM CAY, 50μM LALi, CAY+LALi for 24h or 5µM oligomycin (Oligo, negative control) for 1h. Graphed data represent the mean ± SD for four independent biological replicates. **G-H)** Representative examples of DAN-G cells transfected with cyto-iATPSnFR1.0 and TKS5-mCherry and analyzed by live cell imaging. At t=0, DMSO **(G)** or 50μM LALi **(H)** spiked in, and cells were imaged over 590min. Boxes indicate magnified regions. Quantitation represents cyto-iATPSnFR1.0 mean gray intensity at TKS5 positive puncta per cell at time 0 and 590min. Each line represents the invadopodia fluorescence intensity of one cell. Data represents three independent biological replicates. p values calculated using unpaired Student’s t test for **(A-F)** and paired Student’s t test for **(G-H)**. *p < 0.05, **p < 0.01, ***p < 0.001 ****p < 0.0001, ns = not statistically significant. Scale bar = 10μm; magnified images scale bar = 2µm.

Following the changes in oxygen consumption upon inhibition of lipase activity, we next investigated the levels of reactive oxygen species (ROS) in the PDAC cells. However, we observed only modest effects on intracellular and mitochondrial ROS upon LAListat1 treatment, as measured by H2DCFDA and MitoSOX Red respectively (Figure S5F-I), and the effects varied by cell line. While there may be some contributions to redox regulation by lipid droplet catabolism, this did not appear to be the mechanism by which lipid droplet dynamics contribute to tumor cell invasion.

We next investigated the contributions of lysosomal and cytosolic lipid catabolism to cellular energetics by measuring levels of ATP using an ATP Detection Assay kit. At the whole cell level, inhibiting lipase activity decreased total ATP levels, though effects were highly variable (Figure 5E-F). Inferring ATP production from the Seahorse metabolic assays indicated a clearer inhibition of ATP production upon inhibition of LAL or HSL, with combined treatment showing an additive inhibition (Fig 5D).

As our data suggests, there is spatial regulation of lysosomal lipid droplet catabolism during invasion (Figure 4), we investigated if localized ATP production near invadopodia might also be impacted by lipophagy. To this end, live cell imaging was performed using cells transfected with TKS5-mCherry to label invadopodia and the cytoiATPSnFR1.0 fluorescent ATP biosensor, for which fluorescence intensity is proportional to ATP abundance (*39*). The fluorescence intensity of the ATP biosensor was specifically measured at invadopodia and for the whole cell. The biosensor fluorescence intensity in the whole cell and at invadopodia was stable throughout 590 minutes in cells treated with DMSO (Figure 5G, Figure S5J-K, Supplemental Movie 7-9). Total cell fluorescence indicating ATP levels was only modestly affected in LAListat1 treated cells (Figure S5J, S5L). In marked contrast, after spiking in LAListat1, we observed a substantial decrease in the intensity of the ATP probe at invadopodia in as little as two hours that persisted upon overnight imaging (Figure 5H, Supplemental Movie 10-12). Thus, not only do invadopodia localize near lipid droplets, but these data also indicate that ATP levels are actively regulated at invadopodia by a lipophagic process to maintain their function in cancer cells. Together with our data above, this suggests spatial control of lipid droplet catabolism in invasive protrusions could be used to support tumor cell invasion.

### Lipid droplet catabolism maintains phospholipid homeostasis

Our data suggest that LAL and HSL/MGL both promote tumor cell invasion through different steps of the invasive process. While both lysosomal and cytosolic lipases catabolize neutral lipids, the relative contributions of lipophagy and lipolysis are poorly understood. It is unclear if these two catabolic pathways act on the same species of lipids and whether they have the same metabolic or mechanistic consequences. To investigate this question, we performed lipidomics analysis on mKPC and DAN-G cells treated with inhibitors to LAL, HSL/MGL, or both LAL and HSL/MGL. While we hypothesized that there would be distinct lipid pathways or species impacted by inhibiting lysosomal versus cytosolic lipases, we speculated that these would be conserved between the cell models. In both cell lines and under all lipase inhibitor conditions, there was a significant accumulation of triglycerides and cholesterol esters, consistent with defective lipid droplet catabolism of these target species. Notably, there were heterogeneous effects on the cellular lipid profile between cell types and among inhibitors (Figure 6A-B, Table S1).

**Figure 6.**
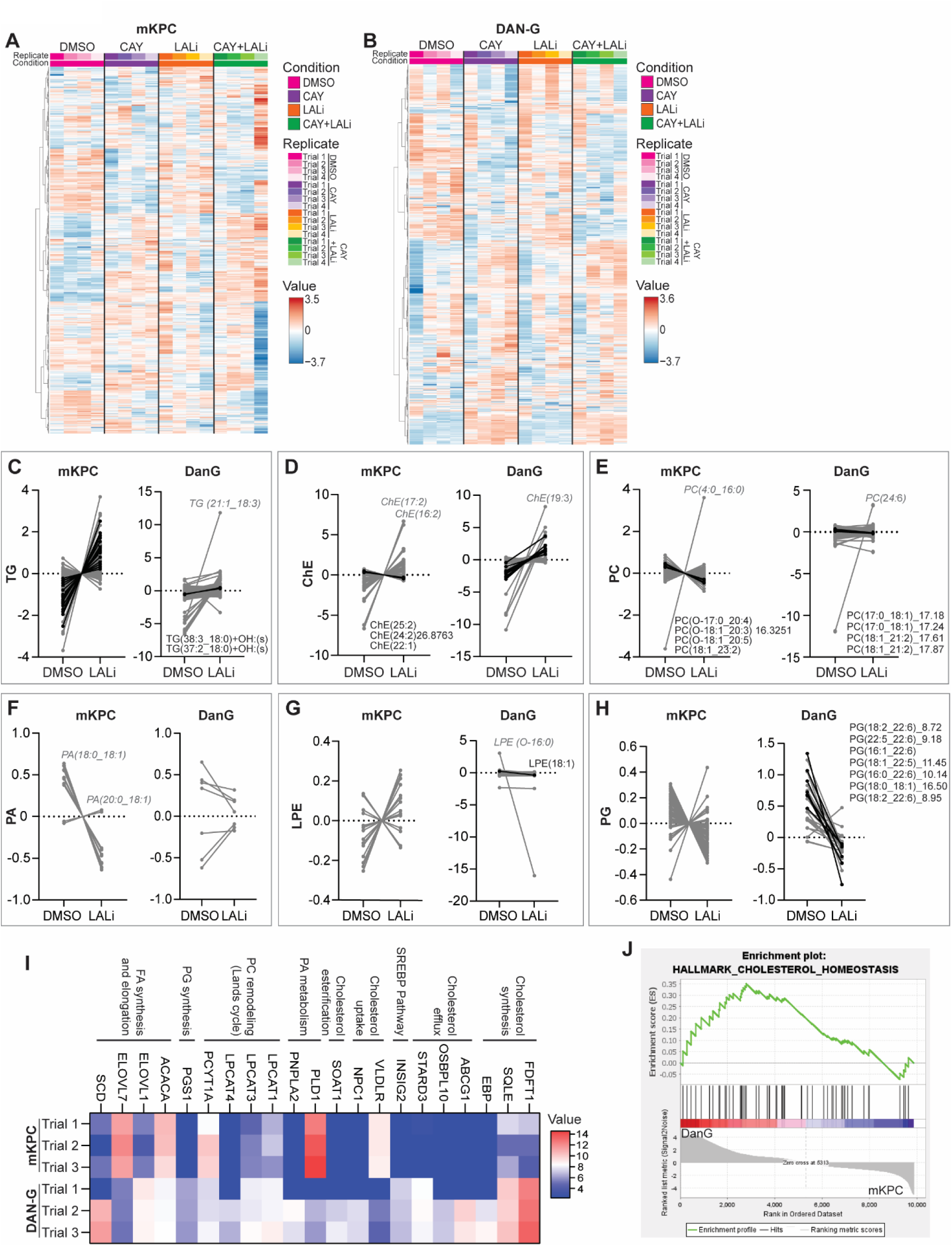
Lysosomal lipolysis modulates membrane lipid composition. **A-B)** Metaboanalyst heatmap of mKPC **(A)** and DAN-G **(B)** cells treated with either DMSO, 10μM CAY, 50μM LALi, or CAY+LALi for 24h. **C-H)** Specific paired lipid species from mKPC and DAN-G cells treated with DMSO or LALi (50μM, 24h) of triglycerides (TG) **(C)**, cholesterol esters (ChE) **(D)**, phosphatidylcholine (PC) **(E)**, phosphatidic acid (PA) **(F)**, lysophosphatidylethanolamine (LPE) **(G)**, and phosphatidylglycerol (PG) **(H)**. Black lines: statistically significant differences. Select individual species are noted. **I)** Heatmap of mKPC and DAN-G RNAseq significantly different gene hits from relevant pathways. **J)** Enrichment plots for DAN-G vs. mKPC cells of MSigDb Hallmark gene set collections for Hallmark cholesterol homeostasis pathway (NES=1.16, Nominal p-value= 0.31, FDR q-value= 0.49). Data represents four independent biological replicates.

We were particularly interested in the smaller subset of lipids impacted by LAL inhibition, as LAL activity had a more potent effect on invadopodial biology than the cytosolic lipases (Figure S6A-B). LAListat1 treatment led to an accumulation of multiple species of triglycerides (TG) and one diglyceride (DG) in mKPC cells, while the DAN-G cells only had a significant accumulation of two TG species (Figure 6C, Figure S6A-C). Instead, in DAN-G cells, LAListat1 induced a significant accumulation of cholesterol esters, particularly with unsaturated fatty acid side chains (Figure 6D, Figure S6B, Table S1). Broadly, alterations of distinct lipid species indicate metabolic variability between cell models.

In addition to the predicted accumulation of neutral lipid species, there were also changes in phospholipid balance upon inhibition of lysosomal versus cytoplasmic lipases. Overall amounts of lipid classes were not globally affected, which indicates specificity towards certain species rather than blockade of metabolic pathways. In both mKPC and DAN-G cells, treatment with LAListat1 increased select forms of phosphatidylcholine (PC), particularly PCs containing unsaturated lipids (Figure 6E, Figure S6A-B). We were particularly interested in which lipid species were deficient in the presence of lipase inhibition, suggesting defective flux from stored neutral lipids. In mKPC cells, we observed decreased phosphatidic acid (PA) and phosphatidylserine (PS) species following LAListat1 treatment (Figure 6F, Figure S6D). In the DAN-G cells there was a notable decrease in phosphatidylglycerol (PG) species, particularly those containing 22:6 (DHA) (Figure 6H, Figure S6B, Table S1).

To identify metabolic pathways impacted by cytosolic versus lysosomal lipases between the cell lines, we performed pathway analysis through BioPAN on LIPID MAPS (Figure S6E-J) (*40*). Inhibition of LAL consistently resulted in imbalance of phosphatidylethanolamine (PE) flux (Figure S6H-I). DAN-G cells in particular showed defects on PE-dependent LPE production, whereas there is predicted to be a deficit in PE-derived PS in mKPC cells in response to lipase inhibition (Figure 6G). Thus, lipid droplet catabolism likely feeds into production or regulation of PE, potentially from fatty acids derived from DG/TGs or from lipid droplet membranes.

The substantial differences in lipid profile between the DAN-G and mKPC cells treated with lipase inhibitors led us to speculate that these two cell lines have different metabolic wiring caused by differences in gene expression programs. We therefore performed RNA sequencing on mKPC and DAN-G and focused on statistically different gene expression in cholesterol regulation and phospholipid remodeling pathways (Figure 6I, Table S2). In line with our previous observations in DAN-G cells, Phosphatidylglycerophosphate Synthase 1 (PGS1) is increased in comparison to mKPC cells, which may explain the dependence upon specific PG species in the lipidomics data set (Figure 6H). We also observed an increase in cholesterol synthesis, efflux and transport in DAN-G cells compared to mKPC cells (Figure 6I), which correlates with a strong dependence on cholesterol and the significant impact of LALi on cholesterol esters in DAN-G cells (Figure 6D). Gene set enrichment analysis suggested that DAN-G cells may show signatures consistent with cholesterol homeostasis, oxidative phosphorylation, and fatty acid metabolism (Figure 6J, Figure S6K-L). Critically, even though the underlying metabolic programs and biochemical changes resulting from lipase inhibitors were different between cell models, lipase inhibition profoundly inhibited transwell invasion in both settings. Thus, lipase targeting may broadly impact tumor cell invasion across subtypes and even in the context of tumor cell heterogeneity.

### Lipid droplet catabolism governs plasma membrane tension and cholesterol content at invadopodia

While the specific lipid species controlled by lipophagy exhibited variability in the DAN-G versus mKPC cells, the common theme of alterations in phospholipid species or fatty acid changes suggested a potential functional impact on membrane fluidity, dynamics, or tension. Aberrations in membrane lipids could impair cell invasion by impacting formation and curvature of nascent invasive protrusions or by altering localized signaling. To test the impact of LAL-dependent membrane remodeling on cell membrane tension, we labeled cells with the Flipper TR probe, which indicates membrane tension *in vitro* by measuring fluorescence lifetime imaging (*41, 42*). After treatment with LAListat1 for 24 hours, there was an increase in cell membrane tension in mKPC cells, though not in DAN-G cells (Figure 7A-B). These data indicate that lipid droplet catabolism is used, at least in part, to maintain membrane lipid pools and thus normal membrane tension.

**Figure 7.**
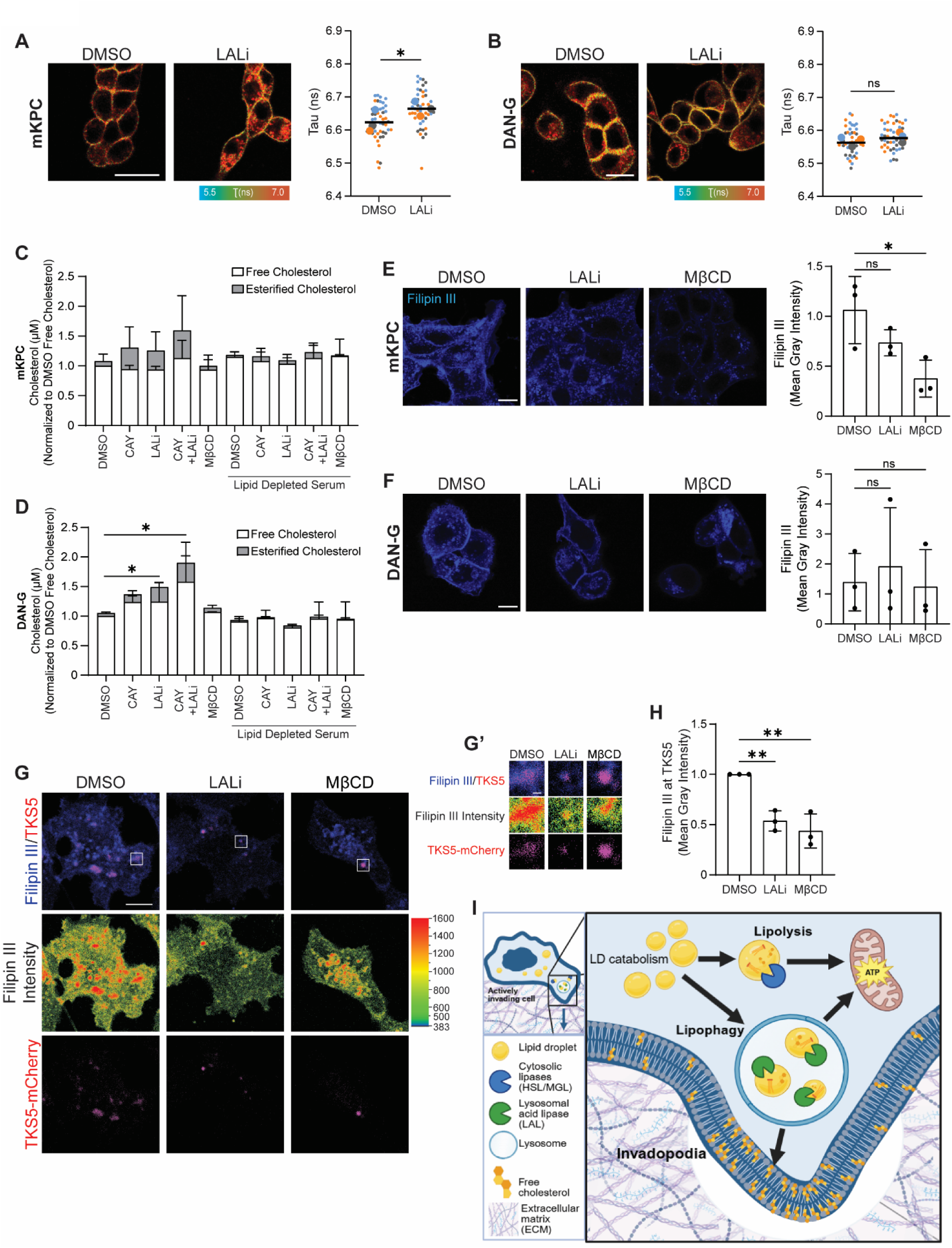
Lysosomal lipolysis modulates membrane cholesterol levels and membrane tension. **A-B)** Fluorescent images of mKPC **(A)** and DAN-G **(B)** cells treated with DMSO or LAListat1 (50 µM, 24h) with Flipper TR Probe. Quantitation of membrane tension based on fluorescence lifetime imaging (Tau). Representative images and quantitation for three independent biological replicates. Data points represent individual cells color coded by biological replicate, and statistics calculated on the mean values of three biological replicates. p values calculated using a paired Student’s t test. Scale bar = 20µm. **C-D)** Free and esterified cholesterol measurements of mKPC **(C)** and DAN-G **(D)** cells treated with DMSO, 10µM CAY, 50 µM LALi, CAY+LALi, or 0.5mM methyl-β-cyclodextrin (MβCD, negative control) under full serum media or media with lipid depleted serum. Data represents the mean ± SD for three independent biological replicates, with cholesterol concentrations below the limit of detection of the kit replaced with the value of zero. Statistical significance was calculated for esterified cholesterol levels. **E-F)** mKPC **(E)** and DAN-G **(F)** cells treated with DMSO, 50µM LALi, or MβCD (5mM 2h for mKPC, 3.5mM 2h for DAN-G) and stained for free and membrane-associated cholesterol with Filipin III (blue). Filipin III average mean gray intensity per cell was measured for three biological replicates. n=10 fields per condition for each replicate. Scale bar = 10µm. **G)** Fluorescent images of DAN-G cells transfected with TKS5mCherry and stained with Filipin III, treated with DMSO or LAListat1 (50 µM, 24h) or MβCD (3.5mM, 2h). Scale bar = 10μm; magnified images **(G’)** scale bar = 1µM. **H)** Quantitation represents Filipin III mean gray intensity at TKS5 positive puncta. Data represents three independent biological replicates. p values were calculated using unpaired Student’s t test. *p < 0.05, **p < 0.01, ns=not significant. **I)** Graphical abstract of lysosomal lipase promoting pancreatic cancer cell invasion through controlling local energetics and lipid remodeling at invadopodia.

Sterols in the plasma membrane, including free cholesterol, can greatly impact cell membrane tension (*43–45*). Since we observed inhibition of cytosolic or lysosomal lipases led to accumulations of cholesterol esters (Figure 6A-B, D, Figure S6A-B), we next measured free and esterified cholesterol. Consistent with the lipidomics analysis, inhibition of LAL or HSL/MGL increased the amount of cellular cholesterol esters in both mKPC and DAN-G cells (Figure 7C-D). While we originally predicted that this would come at the expense of free cholesterol due to impaired flux from cholesterol esters. Instead, lipase inhibition led to a marked increase in total cellular cholesterol. We hypothesized that there may be compensatory upregulation of cholesterol from the culture media following lipase inhibition. To this end, we cultured cells in lipid-depleted serum while in the presence of the lipase inhibitors. Indeed, we found that this completely prevented the increase in cellular cholesterol induced by inhibition of HSL/MGL or LAL (Figure 7C-D), demonstrating the importance of cholesterol uptake to balance lipase inhibition.

We hypothesized whole-cell measurements of free cholesterol or membrane tension might not capture cholesterol levels or distribution specifically in the plasma membrane. We therefore investigated the contributions of LAL-mediated cholesterol metabolism towards invasive cell biology by staining DAN-G and mKPC cells with Filipin III to detect free cholesterol (*46, 47*). As predicted, DMSO-treated cells showed robust filipin-labeled cholesterol in the plasma membrane, intracellular membranes, and accumulation in intracellular compartments, likely lysosomes or endosomes (Figure 7E-F). LAListat1 treatment decreased the amount of vesicular membrane cholesterol, consistent with disrupted lysosomal hydrolysis of cholesterol esters, but did not significantly decrease in overall fluorescence intensity (Figure 7E-F). Membrane cholesterol is a major component of invadopodia, and regulation of membrane fluidity and signaling is essential for these invasive protrusions (*48*). Further, as our data implicate spatial control of lipophagy at invadopodia, we measured the impact of LAL inhibition on membrane cholesterol specifically at invadopodia. Cells were transfected with TKS5-mCherry to label invadopodia, and the fluorescence intensity of Filipin III was measured exclusively at TKS5-mCherry puncta (Figure 7G-H). Critically, in cells treated with LAListat1, there was a substantial 50% decrease in free cholesterol levels specifically at invadopodia compared to DMSO treatment (Figure 7H). This indicates that lysosomal lipolysis via LAL is important in shaping the plasma membrane composition at invadopodia, specifically free cholesterol levels, supporting invadopodia formation and stabilization.

## Discussion

In this study we present evidence for lipid droplet catabolism in the early steps of cancer cell invasion and demonstrate both shared and unique roles for cytosolic versus lysosomal lipases in lipid metabolism and invasive migration (Figure 7I). Expression and activity of both cytosolic HSL and LAL promote invasion by pancreatic cancer cells. However, cytosolic HSL supports 2D migration, whereas LAL had a stronger effect on invadopodia biology and ECM degradation. While both HSL and LAL supported mitochondrial oxidative metabolism, lipidomics revealed distinct lipid substrates and products resulting from HSL versus LAL, with the precise lipid species specific to the cell type. Further, localized lipid droplet distribution and catabolism appear essential for invasive cell biology, regulating both localized energetics and membrane lipids specifically at invadopodia sites. Thus, we propose that lysosomal lipid droplet catabolism supports invadopodia formation through altering both membrane lipid composition and ATP levels.

While a metabolic shift towards glycolysis has been widely described to support tumor growth, increasing evidence suggests oxidative phosphorylation is a major driver of metastatic invasion (*4, 49–51*). Indeed, PDAC cells undergo a metabolic shift towards oxidative phosphorylation during migration and invasion, which required both lipase activity and fatty acid oxidation (*14, 52–54*). Inhibition of either HSL or LAL decreased basal and maximal mitochondrial respiration, as well as ATP levels, coinciding with defects in tumor cell invasion, but not proliferation or viability *in vitro* (Figure 5). We interpret that high levels of ATP production are required to sustain the ATP-intensive process of actin remodeling, which is regarded as one of the most energy rigorous cellular processes (*2, 55*). G-actin monomers conjugated to ATP are incorporated into the growing barbed end of actin filaments, whereas ATP hydrolysis to ADP along the F-actin filament contributes to actin severing and turnover (*56*). Abundant and efficient ATP production are therefore needed to support invasion, which relies on extensive actin cytoskeletal-membrane dynamics. Mitochondrial beta oxidation of fatty acids generates the reducing species needed to drive the electron transport chain, and thus represents a robust mechanism of ATP production.

Furthermore, to fuel global mitochondrial metabolism and ATP production, our data indicates that lipid droplet localization and spatial regulation of lipophagy are drivers of tumor cell invasion, likely through local production of ATP (Figure 4-5). Mislocalizing lipid droplets away from the periphery led to a significant decrease in PDAC invasion (Figure 4). A subset of lipid droplets have been shown at the plasma membrane in *Drosophila* development (*57*), and de novo lipogenesis supports invasive protrusions and prenylation in *C. elegans* (*58*). Another paper observed inhibition of tumor progression with forced clustering of lipid droplets to disrupt organelle contacts (*59*). These data put the spatial regulation of lipid droplets in the context of mammalian cell and invasion. Similarly, mitochondria have been shown to localize at the invasive front in both mammalian cells and *C.elegans*, supporting local ATP production (*2, 3, 5, 60, 61*). We also observed the localized generation of ATP at invadopodia in a manner dependent upon lysosomal acid lipase activity, indicating that not only are localized energetics needed for cancer cells to invade, but that localized lipid droplets also play a crucial role.

In addition to energy production, we speculated that biological differences between cytosolic and lysosomal lipid metabolism may be due to differences in neutral lipid substrates or in the fate of the resulting fatty acid products. Indeed, lipidomics data indicate deficits in specific membrane lipids upon inhibition of lipolysis and lipophagy. Inhibition of cytosolic or lysosomal lipolysis both led to a marked accumulation of cholesterol esters (Figure 6). There were also significant and cell-specific defects in production of specific phospholipid species, including reduced DHA-containing PGs in DAN-G cells and alterations in specific species of PC and PE. It is intriguing to speculate that localized liberation of cholesterol or fatty acids for conversion to membrane phospholipids remodels the local membrane environment, supporting membrane fluidity and signaling that contribute to tumor cell invasion. Invadopodia membranes are rich in saturated phospholipids, cholesterol, and specific signaling lipids such as the phosphoinositides PI(4,5)P_2_ and PI(3,4,5)P_3_ (*48, 62–65*). Lipid droplets specifically contain the phospholipids and cholesterol needed to alter the lipid membrane composition and support membrane fluidity. PC 34:1 has been implicated in invadopodia formation (*66*), and numerous phospholipids including PG, PA and PS and cholesterol have been implicated in the formation of lipid rafts, which are essential for invadopodia formation (*48, 67*). Saturated lipids have also been linked to stiffer cell membranes and decreased invadopodia formation (*66, 67*). We infer that lysosomal lipid droplet catabolism liberates specific fatty acids and cholesterol that could subsequently be incorporated into the plasma membrane to support invadopodia formation. Cell-specific changes in lipids led us to speculate that the production of specific lipid species resulting from lipophagy or lipolysis likely depends on the metabolic, genetic, or epigenetic differences in specific cancer cells that impact both lipid metabolic pathways as well as invasive potential and mode of invasive migration. However, the net consequence of altered membrane tension, fluidity, or local membrane composition could impact migratory potential.

Previously, we found that supplementation with exogenous oleic acid promotes invasion through catabolism by the cytosolic lipase HSL (*14*). Here, we demonstrate that the pro-invasive role of lysosomal lipid catabolism via LAL was restricted to basal media conditions, but was lost upon overloading with exogenous oleic acid. Oleic acid has been used in previous studies to replicate high fat diets in obese patients (*68*), which have been linked to increased PDAC tumor progression by changing the metabolic landscape in the PDAC cells (*69*). This monounsaturated fatty acid can function as an activator in mouse models of PDAC to increase cancer progression (*70*). These findings are consistent with our observations that it is not solely the amount of fat or amount of lipid droplets present, but the specific type of fat that can modulate tumor progression. We did not observe an anti-invasive effect of LAL inhibition in the pancreatic cancer cells upon oleate overload. This suggests that lipid droplet fatty acid content is important for dictating the metabolic and functional consequences of lipid catabolism, and biasing cells towards a specific fatty acid species may interfere with the role of distinct lipid metabolic pathways. Related to the source of fatty acids, LAL is implicated in hydrolysis of triglycerides and cholesterol esters both liberated from storage in lipid droplets as well as internalized as lipoproteins via endocytosis (*71*). The source and means of lysosomal accumulation of these neutral lipids could also drive specificity of lipid substrates or products with biological consequences.

Notably, inhibition of HSL or LAL had no effect on cell viability *in vitro*, similar to prior studies (*14, 21*). As these experiments were conducted in the presence of high glucose, glutamine, and serum, it is possible that any potential pro-survival effects of lipolysis are minimized due to abundance of alternative fuel sources. Intriguingly, overexpression of HSL, which also did not affect viability or proliferation *in vitro*, had a marked effect on tumor growth *in vivo* (*14*). This suggests that the role of HSL may be even more profound in the nutrient-depleted pancreatic tumor microenvironment. On the other hand, altered lipid metabolism may exaggerate or restrict interactions with other cellular components of the complex tumor microenvironment. Future studies will articulate more carefully the relative roles of cytosolic and lysosomal lipolysis in tumor growth and metastasis *in vivo*.

Together, these data demonstrate the role of lipid droplet turnover via both cytosolic HSL and lysosomal acid lipase in PDAC cells, supporting the production of energy and the membrane lipid building blocks needed for invasion.

## Materials and Methods

### Cell Culture

DAN-G, mKPC T42D (a gift from Dr. David Tuveson,(*72*)) and PANC-1 (American Type Culture Collection (ATCC)) cell lines were cultured in DMEM (Corning) containing 10% FBS (Corning, 35-010-CV) and 1% Penicillin/Streptomycin (Gibco, 15140-122). Cells were maintained at 37°C in a 5% CO_2_ incubator. BxPC-3 (ATCC) cells were cultured using RPMI1640 (Corning) containing 10% FBS and Pen/Strep. Cell lines were checked for Mycoplasma using MycoAlert Mycoplasma Detection Kit (Lonza, LT07-318) or by nuclei staining using Hoechst 33342 (Invitrogen, H3570). 50 µM LAListat1 (23891) and 10 µM CAY10499 (10007875) were from Cayman Chemical. For oleic acid (OA) loading, 11.9mM OA stock was made with FAF-BSA (Sigma-Aldrich, A6003-10G), 0.1M Tris and diluted with sterile water. The solution was filtered through a 0.22µm syringe filter (Millex, SLGVR33RS) and allowed to rotate for 10min. Oleic acid (Sigma Aldrich, O1383-1G) was added to the solution and incubated overnight at 4°C rotating and filtered with 0.22µm syringe filter before use, stored at 4°C. 0.2mM OA was used for each 24h treatment.

### Transfections

Lipofectamine 2000 (Invitrogen, 11668027) was used for transfection of TKS5-mCherry (*73*), pcDNA3.1(+)hsaLIPA-FLAG (GenScript, OHu23443) and pcDNA3.1 FLAG (a gift from Dr. Daniel Billadeau). Lipofectamine 3000 (Invitrogen, L3000008) was used for transfection of pEGFP-N1-LAMP1 (*74*), TKS5-GFP (*73*), RFP-eGFP-PLIN2 (*21*), cyto-iATPSnFR1.0 (Addgene, plasmid #102550, (*39*)), or TKS5-mCherry. For knockdowns, the *LIPE* gene encoding HSL was targeted with RNAi against mouse *Lipe* (Dharmacon # M-040205-01-0010) or human *LIPE* (Dharmacon custom 5’-GCAGCCUGAUAAAGUCCAATT-3’), and the *LIPA* gene encoding LAL was targeted with RNAi against mouse *Lipa* (Dharmacon # M-048418-01-0005) or human *LIPA* (Dharmacon # L-004043-00-0005) with Lipofectamine RNAiMAX (Invitrogen, 13778030). GFP-KIN14-FRB and RFP-PEX-FKBP were kindly provided by Dr. Lucas Kapitein (*38*). PLIN2 was amplified from RFP-GFP-PLIN2 (*21*) to replace PEX using Gibson cloning and using NheI (New England Biolabs, R0131) to linearize the RFP-PEX-FKBP vector. Primers to amplify PLIN2 were: Forward: 5’-CCCTTCTCTGTTCCTCCGCAGCCCCCAAGCTAGCATGGCATCCGTTGCAGTTGATCCACAACCGAGTGTGG-3’; Reverse: 5’-CCCGACCCATTTGCTGTCCACCAGTCATGCTAGCATGAGTTTTATGCTCAGATCGCTGGGTCTCCTGGC-3’.

### CRISPR KO cell lines

LAL was knocked out in mKPC T4 2D cells using the CRISPR/Cas9 system. CHOP CHOP CHOP (*75*) was used to design gRNA target sequence 5’-ACCGAGATAATCATGCGCTGGGG -3’ to target Exon 3 of *LIPA* in the *Mus musculus* (mm10/GRCm38) with the genomic coordinates of chr19:34523517. The gRNA was cloned into lentiCRISPR v2 (Addgene plasmid #52961, gift from Dr. Feng Zhang,(*76*)) as previously described (*77*) with modifications. LentiCRISPR v2 plasmid was digested with BsmBI (New England Biolabs, R0739S), and then treated with Quick CIP (NEB, M0525S) prior to electrophoresis and gel purification (QIAquick PCR and Gel Cleanup Kit (Qiagen, 28506). To phosphorylate and anneal the pair of oligos, 100µM of Oligo 1 (5’-CACCGACCGAGATAATCATGCGCTG-3’) and Oligo 2 (5’-AAACCAGCGCATGATTATCTCGGTC-3’) were incubated with T4 Ligation Buffer (New England Biolabs, B0202S) and T4 Polynucleotide Kinase (New England Biolabs, M0201S), and the ligation was set up at room temperature following “Target Guide Sequence Cloning Protocol” from GeCKO (*77*) and transformed into NEB Stable Competent E. coli (High Efficiency) (NEB, C30401). LentiCRISPR v2 LAL gRNA or control were transfected into mKPC cells. Cells underwent puromycin selection at 1μg/mL for 48h post transfection and single clones were selected and expanded.

To check LAL KO, nuclei were isolated with gentle lysis buffer (150mM NaCl, 20mM EDTA pH 8.0, 50mM Tris-HCl pH 7.5, 0.5% NP-40, 1% Triton X-100, 20mM NaF) and a nuclear lysis buffer was applied (150mM NaCl, 20mM EDTA pH 8, 50mM Tris-HSL pH 8, 1% NP-40, 20mM NaF, 0.5% Na deoxycholate, 0.1% SDS) to isolate the genomic DNA. RNAse A was added (100µg/mL final concentration) for each sample and incubated for 30min at 37°C at 700rpm. Proteinase K was added and allowed to incubate rotating for 16h at 55°C following with the addition of phenol/choloroform/isoamylic alcohol (25:24:1). The samples were vortexed for 30s and spun down at full speed for 2min. The top aqueous phase was taken, followed by the addition of 3M sodium acetate pH 5.5 (10%) and glycogen (1%). An ethanol precipitation followed with overnight incubation at −20°C, with cells resuspended in Tris-EDTA Buffer.

Following DNA isolation, a nested PCR was performed with the 2x Platinum SuperFi II Green PCR Master Mix (Invitrogen, 12369010) following the manufacturer’s protocol. Outer forward primer (5’-GATGTTAAATGAAGGTATTATACAGCAAGA-3’) and outer reverse primer (3’-TTCTTCTATTAGTTTCATGCCTTTCAAATG-3’) were used with 2.5min initial denaturation at 98°C, 35 cycles of 10s denaturation at 98°C, 10s annealing at 60°C, 30s extension at 72°C, followed with a final extension at 72°C for 5min. PCR cleanup was done using KAPA Pure Beads (Roche Applied Science, 07983271001) at 0.7x, vortexed and incubated at room temperature for 5min. The PCR product was spun down, placed on the outer primer PCR product was washed with 80% ethanol and incubated for 30s at room temperature. The supernatant was removed, beads were allowed to dry for 3min, product with beads was spun down again and added to magnet to ensure all supernatant was removed. Cells were incubated in Tris-EDTA buffer for 2min at room temperature, the tube was placed on the magnet, and the supernatant was collected into a new tube. Purified product was run through a second PCR reaction with the inner forward primer (5’-AAATGGAATGTGACCTTAGAACCC-3’) and inner reverse primer (5’-ACAAGAGGCCAAATTTGATACACT-3’) in a second reaction following the 2x Platinum SuperFi II Green PCR Master Mix user guide again with the initial denaturation 30s instead of 2.5min. The same procedure was followed with the KAPA Pure Bead extraction, with a concentration of 1.0x instead of 0.7x for the beads. PCR purified product was Sanger sequenced by GENEeneWIZ (Azenta Life Sciences). After sequences were analyzed using Decodr, CRISPR editing yielded two highly efficient LAL deficient lines. LAL KO Clone 1 revealed frameshift mutations with indels −17, −13 and −5bp, indicating a KO. LAL KO Clone 2 has frameshift mutations at −94bp, −10bp and −232bp indels, as well as a 29.4% contribution of an in-frame deletion at −24bp (Table S3).

### Immunoblotting

For Western blotting, cells were lysed using NP-40 lysis buffer (137mM NaCl, 20mM Tris-Cl, 10% glycerol, 1% IGEPAL, 2mM EDTA (pH8.0)) with complete protease inhibitor cocktail (Roche, 11873580001). BCA protein assay was used to calculate protein concentration and load equal amounts of protein in each well of a SDS-PAGE gel (Bio-Rad) including the molecular weight marker Precision Plus Protein Blue Prestained Standard (Bio-Rad, 1610373). The gel was transferred to a PVDF membrane for 30min using a semidry transfer apparatus and blocked in 5% BSA in TBS. Blots were probed for 2h or overnight with primary antibodies diluted in TBST + 5% BSA. For primary antibodies, HSL (Cell Signaling Technology, 4107S), Actin (Sigma, A2066) and GAPDH (Cell Signaling Technology, 2118S) were used. Secondary antibodies were Goat anti-rabbit horseradish peroxidase (Fisher Scientific, G21234). Western blots were visualized using enhanced chemiluminescence (Thermo Fisher Scientific, 34580), imaged on an Azure 500 Biosystems Imaging System, and band intensity was analyzed using AzureSpot Pro. Source data (uncropped western blots) are deposited at Mendeley Data at DOI 10.17632/hnsfnvgjhv.1.

### qPCR mRNA Analysis

Knockdown of LAL was validated using qPCR. RNA was isolated using PureLink^TM^ RNA Mini Kit (Thermo Fisher Scientific, 12183018A). Isolated RNA was then reverse transcribed with iScript Reverse Transcription Supermix (BioRad, 1708840). LAL mRNA expression was quantitated with PowerTrak^TM^ Master Mix (Thermo Fisher Scientific, A46109) according to manufacturer’s protocol using the QuantStudio3 (Applied Biosystems). 10ng of cDNA was used per sample and primers were: human *LIPA*: Fwd 5’-TCGCCTTCTGTACTAGCCCT-3’, Rev 5’-GCCTGGCTCCAGTGTAACAT-3’ or mouse *Lipa*: Fwd 5’-TGCCCACGGGAACTGTATC-3’, Rev 5’-ATCCCCAGCGCATGATTATCT-3’. GAPDH primers were used for normalization: human Fwd 5’-GGGAGCCAAAAGGGTCATCA-3’, human Rev 5’-GCATGGACTGTGGTCATGAGT-3’, mouse Fwd 5’-ATGGTGAAGGTCGGTGTGAA-3’, mouse Rev 5’-GGGATTACGGGATGGGTCTG-3’. Relative LAL mRNA expression was measured by normalizing LAL to GAPDH using the ΔΔCt method.

### Cell Viability

PANC-1, DAN-G, or BxPC-3 cells (1×10^4^) and mKPC (5×10^3^) cells were plated in triplicate for each drug treatment in a 96-well plate. After 24h, cells were treated with either Dimethyl Sulfoxide (DMSO, Sigma, D2438), 10μM CAY10499, 50μM LAListat1, a combination of 10μM CAY10499 and 50μM LAListat1, or 5μM Staurosporine as a control to induce cell death. Cells were incubated with drugs for 24h before adding 10μL of the Cell Counting Kit-8 (CCK8) reagent (MedChem Express, HY-K0301). After a 1h incubation with the CCK8 reagent, absorbance at 450nm was read on a plate reader (Tecan Infinite 200 Pro). Data were normalized to DMSO set at 100% viability.

### Immunofluorescence Staining

Cells were fixed with formaldehyde solution (3% formaldehyde, 0.1M PIPES (pH 6.9), 1mM EGTA, 3mM MgSO4) for 20min at 37°C. For lipid droplet staining, cells were fixed and then permeabilized and lipids extracted using 60% isopropanol for 30s, stained with 60% Oil Red O solution (5mg/mL Oil Red O (Sigma, O0625) in isopropanol, then diluted in dH_2_O for 60% solution) for 1min and 45s and washed for 30s again with 60% isopropanol. If further antibody staining was required, cells were permeabilized using 0.1% Triton X-100 (Sigma, 93443) in dPBS for 2min, washed immediately afterwards with dPBS, and incubated with blocking buffer (5% goat serum, 5% glycerol, 0.04% sodium-azide (pH 7.2)) for 1h at 37°C. The primary antibodies used were anti-TKS5 (Sigma Aldrich, MABT336) and anti-FLAG (Sigma-Aldrich, F3165). Secondary antibodies were anti-mouse Alexa Fluor 405S (Biotium, ABIN6198068) and anti-mouse Alexa Fluor 568 (Invitrogen, A11004). For Actin staining, Phalloidin-Tetramethylrhodamine B isothiocyanate (TRITC) (Sigma, P1951) was used at 1:300.

Nuclei were stained using Hoechst 33342 (Invitrogen, H3570). Coverslips were mounted on imaging slides using ProLong Gold (Invitrogen, P36934). Fluorescently stained cells were imaged on a 40x oil lens on a Zeiss780 Confocal microscope or 63x oil lens on a Zeiss LSM980 Confocal microscope and images were acquired using Zeiss Zen software. For lipid droplet and invadopodia quantitation, after uploading raw czi files to ImageJ, Gaussian blur was applied to the images to remove background noise. LD area and LD count was quantitated by applying Auto Threshold and analyzing particles.

### Live Cell Microscopy

Cells were seeded in 35mm glass bottom cell culture dishes (Cell E&G, GBD00004-200). Imaging was performed on Zeiss LSM980 Confocal microscope with a 63x oil objective lens. For live cell video analysis of TKS5-GFP and lipid droplets, transfected cells were stained for lipid droplets using LipidTOX Deep Red (Thermo Fisher Scientific, H34477) and imaged at 1 frame every 15s for DMSO treatment (Trial 1: 180 frames, Trials 2 and 3: 240 frames) and for LAListat1 treatment (All replicates 240 frames).

Interactions between TKS5-GFP and lipid droplets were annotated using ImageJ manually frame by frame. For the PANC1 OA wo movies, alongside LipidTOX Deep Red, Lysotracker Blue DND-22 (Thermo Fisher Scientific, L7525) and MitoTracker Green FM (Thermo Fisher Scientific, M7514) were used. For TKS5-mCherry and cyto-iATPSnFR1.0, cells were imaged at 1 frame every 10min for DMSO treatment (Trial 1: 89 frames, Trial 2: 91 frames, Trial 3: 92 frames) and for LAListat1 treatment (Trial 1: 128 frames, Trial 2: 90 frames, Trial 3: 98 frames) at 10 different locations in each imaging dish. Using ImageJ, TKS5 was thresholded and binary masked, and mean gray intensity measurements of cyto-iATPSnFR1.0 levels were measured under the TKS5 positive areas.

### Migration, Invasion and Degradation

For transwell invasion assays, 6-well transwell chambers with 8µm pores (Corning, 3428) were coated with Matrigel (1mg/mL, serum-free media) (Corning, 356231). 300,000 cells were seeded in the upper chamber in serum-free medium and were allowed to invade for either 24h (mKPC and DAN-G) or 48h (PANC-1) towards media containing 10% FBS in the lower chamber. Filters were cut out and fixed with 3% formaldehyde fixing solution. The nuclei of the cells on the top and bottom of the filter were stained with Hoechst 33342 without scraping and mounted on imaging slides using ProLong Gold (Invitrogen, P36934). Both sides of the filter were imaged and nuclei counted on the top and bottom of the filter using a 40x lens on a Zeiss Observer D1 fluorescence microscope with a Zeiss Arclamp power supply (#910426) and Axiocam 305 monocamera or a 60x oil lens on a Nikon Ti2E fluorescence microscope. The percent invasion was determined by quantitation of invaded cells ÷ (invaded + non-invaded cells) x 100. For overexpression experiments, transwell filters were stained and only the overexpressing cells were scored. For wound healing migration assays, cells were grown to confluence in Ibidi Culture-Insert 2 well system (Ibidi, 80209). For experiments using chemical inhibitors, cells were pretreated 24h prior to the start of migration. The insert was removed carefully, the cells were washed with HBSS, fresh media (with 10% FBS and Pen/Strep) was added with or without chemical inhibitors, and incubated at 37°C to allow cells to migrate for 12h. Migration distances were quantitated using ImageJ by drawing the border on the leading edge of the migrating cells and measuring the area (*78*).For gelatin degradation assays, gelatin coated coverslips were prepared as described previously using 1:8 ratio Oregon Green 488 gelatin from porcine skin (Thermo Fisher Scientific, G13186) and 0.2% unlabeled gelatin (Sigma, G1890). Cells were plated on the gelatin-coated coverslips and allowed to degrade the matrix for 8h (DAN-G) or 24h (BxPC-3). Cells were fixed with 3% formaldehyde solution as above, actin was stained with Phalloidin-Tetramethylrhodamine B isothiocyanate (TRITC) (Sigma, P1951) and nuclei were stained with Hoechst 33342. Images were acquired using Zeiss780 Confocal microscope or 60x oil lens on a Nikon Ti2 fluorescence microscope using LED excitation of 405nm, 480nm, and 640nm. Using ImageJ analysis, the phalloidin stain was used to trace the outline of each cell to measure the total cell area. The green channel (green gelatin) was inverted, adjusted auto local threshold to select all spots of degradation, and measured the area of degradation spots and divided by the total cell area to calculate the percent of cell area degradation.

### Lipid Droplet Redistribution Assays

mKPC or PANC-1 cells were co-transfected with RFP-PLIN2-FKBP and GFP-KIN14-FRB using Lipofectamine 3000. For validation of lipid droplet labeling, cells were plated on coverslips for 48h and then treated with 10nM Rapalog A/C heterodimerizer AP21967 (TaKaRa 635056) for 4h prior to fixation with 3% formaldehyde. Following permeabilization with 0.1% Triton X-100 in dPBS for 10min at room temperature, the cells were stained with AUTODOT (Monodansylpentane, MDH) (100 µM, Abcepta SM1000a) for 15min at 37°C to stain the lipid droplets. After washing with dPBS, the coverslips were mounted on imaging slides using ProLong Gold and imaged one hour later. For transwell assays, the transfected cells were pretreated with 10nM Rapalog A/C heterodimerizer AP21967 for 4h prior to plating on transwells, incubated at 37°C for 24h (mKPC) or 48h (PANC-1), fixed and stained nuclei as described above. Only the transfected cells were counted on the top (non-invaded) and bottom (invaded) of the filter to calculate the percent of cells invaded.

### ROS and ATP Measurements

Reactive oxygen species (ROS) were measured using 2’,7’-dichlorodihydrofluorescein diacetate (H2DCFDA) assay and MitoSOX Red (Thermo Fisher Scientific, M36007). For H2DCFDA, cells were seeded (1×10^3^ mKPC, 2×10^3^ DAN-G, 5×10^3^ PANC-1) in black 96 well microplates with clear bottoms (Corning Falcon, CLS353219) 48h prior to readout. After 1 day of seeding, DMSO, 10μM CAY10499 and/or 50μM LAListat1 was added to each well for 24h and incubated at 37°C. Negative and positive controls were 5mM N-acetylcysteine (NAC) and 1mM H_2_O_2_ respectively, which were dissolved in dPBS and added to the cells for 15min before the readout. Prior to the readouts, H2DCDFA was added to each well at 5µM and incubated for 60min at 37°C. Readouts were completed on the Tecan Infinite M Plex plate reader at 493nm excitation and 522nm emission, 110 Gain, 20μs integration time and 25 flashes with 6 well readout per condition. To measure mitochondrial ROS, DAN-G cells were plated at 70% confluence in 6-well plates for 24h and then treated with 10μM CAY10499 and/or 50μM LAListat1 for 24h. Rotenone was used as a positive control (1µM, 2h, Sigma-Aldrich 557368). After 30min incubation with MitoSOX Red, cells were trypsinized, washed with PBS containing 2% BSA (Sigma-Aldrich, A9647), spun down and resuspended in PBS containing 2% BSA. The gating strategy was FSC-A vs. SSC-A for viable cells and then single cell selection with FSC-A vs. FSC-H, then ran 20,000 cells and collected data with 355-A laser with excitation 355nm and emission 610nm. The cells were processed with LSRFortessa and analyzed with FlowJo software (v11.2) (Tree Star, OR, USA). To assess ATP levels, cells were plated (6×10^3^ mKPC, 12×10^3^ PANC-1) in white bottom 96 well microplates (Corning Falcon, CLS353377) for 24h prior to adding 10μM CAY10499 and/or 50μM LAListat1 for 24h. ATP was measured using the ATP Detection Assay Kit (Cayman Chemical, 700410). Readouts were completed on the Tecan Infinite M Plex plate reader for luminescence without attenuation, 1s integration time.

### Seahorse Assay

Oxygen consumption rate (OCR) (pmol/min) and extracellular acidification rate (ECAR) (mpH/min) were measured using the Agilent Seahorse XFe24 extracellular flux analyzer (Seahorse Biosciences). 8×10^3^ DAN-G cells were seeded for 24h on Agilent Seahorse XF24 cell culture microplates (Agilent, 100777-004), treated with DMSO, 10μM CAY, 50μM LALi, 10μM CAY + 50μM LALi, or no treatment for the following 24h. Seahorse XF Base medium (pH 7.4) supplemented with 10mM glucose, 1mM pyruvate and 2mM glutamine was added onto the cells. Chemical inhibitors were also supplemented in the media to be present throughout the assay. After incubation of the microplate at 37°C with no CO_2_ present for 1h, the Agilent Seahorse XFe24 Extracellular Flux Assay Kit (Agilent, B20925) was used to allow for sequential addition of 1μM Oligomycin (Oligo), 2μM FCCP, and 1μM Antimycin A/ 1μM Rotenone. OCR and ECAR were measured by the Agilent Seahorse XFe24 extracellular flux analyzer. Cells were washed with PBS and frozen at −80°C until BCA Protein Assay (Thermo Scientific, 23225) was used to determine the protein concentration. Three replicate measurements were completed for each condition and normalized to total protein amount. For calculations under specific parameters, the report generator user guide was followed. To calculate Basal Respiration, non-mitochondrial respiration rate was subtracted from the last rate measurement before oligomycin injection. Maximal respiration was calculated by subtracting non-mitochondrial respiration rate from maximal rate measurement after FCCP injection. H+ proton leak was calculated subtracting non-mitochondrial respiration from minimum rate measurement after oligomycin injection. ATP production was calculated by subtracting minimum rate measurements after oligomycin injection from the last rate measurements before oligomycin injection. Spare respiratory capacity was calculated by subtracting basal respiration from maximal respiration. Coupling efficiency was calculated by dividing the ATP production rate to the basal respiration rate and multiplying by 100. All of these calculations were normalized to DMSO.

### Fatty Acid Oxidation Assay

Fatty Acid Oxidation (FAO) activity was measured by monitoring the conversion rate of [U-^13^C]palmitate to [M+2, 4, 6]-labeled ^13^C citrate with GC/MS. In brief, the cells were incubated with 100µM [U-13C] palmitate-BSA conjugate overnight. After the incubation, the cells were quickly rinsed in ice-cold PBS and lysed with 80% methanol. The crude lysates were centrifuged to remove debris. The resulting supernatants were dried with nitrogen gas, dissolved in 75μL dimethylformamide (DMF) then derivatized with 75μL N-Methyl-N-(tert-butyldimethylsilyl)trifluoroacetamide (MTBSTFA) + 1% tertbutyldimetheylchlorosilane (TBDMCS) (Regis). Samples were incubated at room temperature for 30min and then analyzed using an Agilent 7890B GC System coupled to a 5977A mass detector. 3μL of derivatized sample were injected into an Agilent HP-5ms Ultra Inert column, and the GC oven temperature increased at 15°C/min up to 215°C, followed by 5°C/min up to 260°C, and finally at 25°C/min up to 325°C. The MS was operated in split-less mode with electron impact mode at 70 eV. Mass range of 50-700 was analyzed, recorded at 1,562 mass units/second. Data was analyzed using Agilent MassHunter Workstation Analysis and Agilent MSD ChemStation Data Analysis software. IsoPat^2^ software was used to adjust for natural abundance as previously performed (*79*). The extent of isotopic ^13^C labeling in citrate was further divided by percent isotopic enrichment of intracellular [U-^13^C]palmitate to determine the conversion rate of [U-^13^C]palmitate to [M+2, 4, 6]-labeled citrate in cells after normalizing the pool sizes of palmitate and citrate with use of D5-palmitate as an internal standard. Palmitate and citrate were detected by GC-MS as TBDMS derivatives at the following m/z values: palmitate (m/z 313), D5-palmitate (m/z 318), [U-^13^C]-palmitate (m/z 329), and citrate (m/z 459-465).

### Lipidomics

mKPC and DAN-G cells were seeded in 10cm culture dishes for 24h, and then DMSO, 10µM CAY10499 and/or 50µM LAListat1 were added for 24h before isolation at ∼90% confluence. Cells were washed with HBSS, trypsinized and resuspended into DMEM with 10% FBS and Penicillin/Streptomycin. Cells were spun down at 3,000g for 5min and washed with sterile PBS twice. Cell pellets were flash frozen and subsequently processed by the Mayo Clinic Metabolomics Core Facility. Lipids were extracted from plasma or cell lysates with methyl tert-butyl ether, dried down and resuspended in mobile phase prior to analyzing by LC/MS. Data were acquired using an Orbitrap Exploris 120 mass spectrometer, coupled with an UltiMate 3000 Binary Rapid Separation LC System (ThermoFisher, USA). Lipids were separated chromatographically on an Accucore C30 HPLC column (2.6µm, 2.1 x 250mm; ThermoFisher, USA). The analysis was performed in both positive and negative electrospray ionization modes, with full scan coverage from m/z 200-1700 at 120K resolution. Data-dependent acquisition targets the top 4 ions for fragmentation at 30K resolution. Sub-ppm accuracy is maintained throughout the acquisition using ThermoFisher’s Easy-IC internal calibrant.

Data processing: Raw data files were imported into LipidSearch software for annotation and alignment. MS peak areas were integrated, and lipids were identified by comparing MS/MS fragment ions with an internal database. After annotation, peak alignment combines results from both positive and negative ionization modes. All peaks were manually reviewed to prevent false identifications. The R package MetaboanalystR was used for data normalization, differential expression analysis, and visualization. In each mode, metabolites were row-wise normalized to a constant sum (SumNorm), log_2_ transformed, and scaled by mean centering (MeanCenter), replacing 0 or NA with ⅕ of the minimum number in the data set to generate the heatmaps using RStudio and Metaboanalyst). Normalized data were analyzed by multivariate approaches such as Principal Component Analysis (PCA) to reveal data heterogeneity, groupings, outliers and trends. A principal component analysis, heat map and variable importance plot resulting from Partial Least Square discrimination analysis (PLS-DA) were obtained for analysis. Univariate statistical analysis, Student’s unpaired t test between conditions provided were performed with multiple testing correction to identify differentially expressed metabolites between two groups with statistical significance (FDR adjusted p-value <= 0.05 and |fold change| >= 1.5). Lipid pathway analysis was performed using BioPAN through Lipid Maps. After separate conversion though LipidLynxX, lipid species were processed at the annotation level of both sum composition and full structure. Visualization of lipid species reactions were calculated at the lipid subclass level in the suppressed status with p-value of 0.05.

### Bulk RNA sequencing analysis

mKPC and DAN-G cells were isolated at ∼80% confluence from T75 flasks. Cells were washed with PBS, trypsinized and resuspended in DMEM with 10% FBS and Penicillin/Streptomycin. 1.5×10^5^ cells were used per replicate and spun down for 5min at 300xg and washed again with PBS. After the second spin down for 5min, supernatant was aspirated again and cells were resuspended into Zymo 1x DNA/RNA Shield (Zymo Research, R1100). Cells were vortexed for 2min to sheer chromosomal DNA. Three biological replicates were sent per cell line to Plasmidsaurus, each cell line shipped separately at room temperature for RNA sequencing analysis.

RNA sequencing was performed by Plasmidsaurus using Illumina Sequencing Technology with custom analysis and annotation. Plasmidsaurus processing of FastQ generation and demultiplexing was performed by BCL Convert v4.3.6 and fqtk v0.3.1. Comprehensive QC report was generated using MultiQC v1.33 and mapping QC generated by RustQC v0.2.1. Raw sequence reads were aligned appropriate reference genome using STAR aligner v2.7. BAM files sorted using samtools v1.21. Gene-expression quantification was utilized with featureCounts (subread package v2.1.1).

Data processing: mKPC and DAN-G raw counts were used for the analysis in R studio utilizing the Ensembl Gene IDs. The mKPC gene set was converted to the human orthologs through the org.Hs.eg.db annotation database. The dataset was then normalized with DESeq2 median-to-ratios to consider sequencing depth differences. Then we performed variable stabilizing transformation (VST) to take the variability of the gene expression counts into account, followed with principal component analysis and heatmap creation considering cholesterol regulatory and phospholipid remodeling pathways. Gene enrichment analysis (GSEA) (*80, 81*) was performed on DESeq2 dataset (*82*) with 1000 permutations, with the Molecular Signatures Database (MSigDB) with Human Ensembl Gene ID platform (v2026.1.Hs) and with default parameters. RNA sequencing Data are deposited in BioProject (PRJNA1474006).

### Cholesterol Measurement Assays

mKPC (5×10^3^ cells) or DAN-G (1×10^4^ cells) were plated in triplicate for each drug treatment in a 96-well plate. After 24h, media was washed out and replaced with media containing 10% FBS or 10% lipid-depleted FBS (BioWest, S148L). Cells were treated with either DMSO, 10μM CAY10499, 50μM LAListat1, a combination of 10μM CAY10499 and 50μM LAListat1, or 0.5mM Methyl-β-cyclodextrin (MβCD) as a negative control. Cells were incubated for 24h and then the confluency of each well was measured using an IncuCyteS3. Free and esterified cholesterol were then measured using a commercially available Cholesterol/ Cholesterol Ester-Glo^TM^ Assay kit (Promega #J3190) following the manufacture’s protocol. Data were normalized using confluency measurements and normalized to DMSO free cholesterol. Free cholesterol was visualized using Filipin III (Sigma, F4767). mKPC and DAN-G cells were plated at 70% confluence and treated with either DMSO or 50µM LAListat1 for 24h, or MβCD (5mM for mKPC, 3.5mM for DAN-G) for 2h before fixing. After formaldehyde fixation, Filipin III stain was added at 50µg/mL for 1h at room temperature and then mounted on imaging slides with ProLong Gold. Filipin III cells were traced and mean gray intensity was measured. For Filipin III at invadopodia, TKS5-mCherry transfected cells were traced, converted to 8-bit, thresholded, then binary masked. Using the Image calculator, Filipin III channels were multiplied by the TKS5 binary mask and measured for mean gray intensity.

### Membrane Tension Analysis

Plasma membrane tension/packing was assessed in live DAN-G and mKPC cells using fluorescence lifetime imaging and the Flipper-TR probe. Cells grown on glass-bottom multi-well plates were treated with DMSO or 50 µM LAListat1 for 24h. The existing treatment media was then temporarily removed, supplemented with 1 µM Flipper-TR, and returned to the cells for a 10-min incubation at 37°C and 5% CO2, then imaged immediately on an inverted Leica Stellaris 8 equipped for TCSPC-FLIM using 488-nm pulsed excitation and a 600 ± 50 nm emission window. ROIs were drawn along the plasma membrane in a z-plane approximately 2 µm above the coverslip to avoid basal background. Photon-arrival histograms were fitted in LAS X using a double-exponential decay model, and [τlong / amplitude-weighted mean lifetime] was used as the tension-sensitive readout. Data represents 45 cells from three independent biological replicates.

### Statistical Analysis

All microscopy images quantitation was performed using ImageJ and images were processed via Zen software, Nikon Elements software, and Adobe Illustrator. Image acquisition settings were kept consistent with each experiment. All analysis was done on images with equal settings. Representative images shown in figures have individually adjusted brightness levels to most clearly display representative data only in images where fluorescence intensity was not measured. For all microscopy images and immunoblots, linear adjustments in brightness and contrast were applied uniformly across the entire image. Data were analyzed and graphed using GraphPad Prism. Superplots were used to visualize both individual datapoints and statistics on means from independent biological replicates (*83*). Graphs represent the mean +/- SD of at least three biological replicates per condition and statistical significance was calculated using a Student’s t test, unless otherwise indicated. The CCLE gene expression data was analyzed using GEO Expression Omnibus.

## Acknowledgments

We acknowledge the Mayo Clinic Metabolomics Core for the performance and primary analysis of the Lipidomics experiments. We acknowledge Dr. Omar Gutierrez Ruiz, Dr. Kevin Burton, Eugene Krueger, William Taunton, and Yash Saoji for experimental and technical assistance. The model in Figure 7I was created in BioRender. Nooren, R. (2026) https://BioRender.com/vug286v.

## Author Contributions

Conceptualization: RN, CR, JL, MM, GR; Formal Analysis: RN, KJ, MGM, CR, DG, ZOC, MEG, TE; Funding Acquisition: JL, MAM, GR; Investigation: RN, KJ, MGM, CR, DG, ZOC, MEG, TE; Methodology: RN, KJ, MGM, CR, DG, ZOC, MEG, TE, DB, TH, IL Resources: DB, TH, IL, MM, GR; Supervision: GR; Validation: RN, KJ, MGM, CR, DG, ZOC, MEG, TE; Visualization: RN, KJ, MGM, CR, MEG; Writing – Original Draft: RN, GR; Writing – Review & Editing: All Authors.

## Data availability

All data needed to evaluate the conclusions in the paper are present in the paper and/or the Supplementary Materials. Source data for western blots are deposited at Mendeley Data at DOI 10.17632/hnsfnvgjhv.1.

