## Supplemental Figures S1-S6 for "Lysosomal lipid metabolism promotes tumor cell invasion through local energetics and membrane lipid remodeling"

Roseanne E. Nooren *et al.*

**This PDF file includes:**

Figures S1 to S6

**Other Supplementary Materials for this manuscript include the following:**

Movies S1 to S12
Data Tables S1 to S3

Supplemental Figure S1

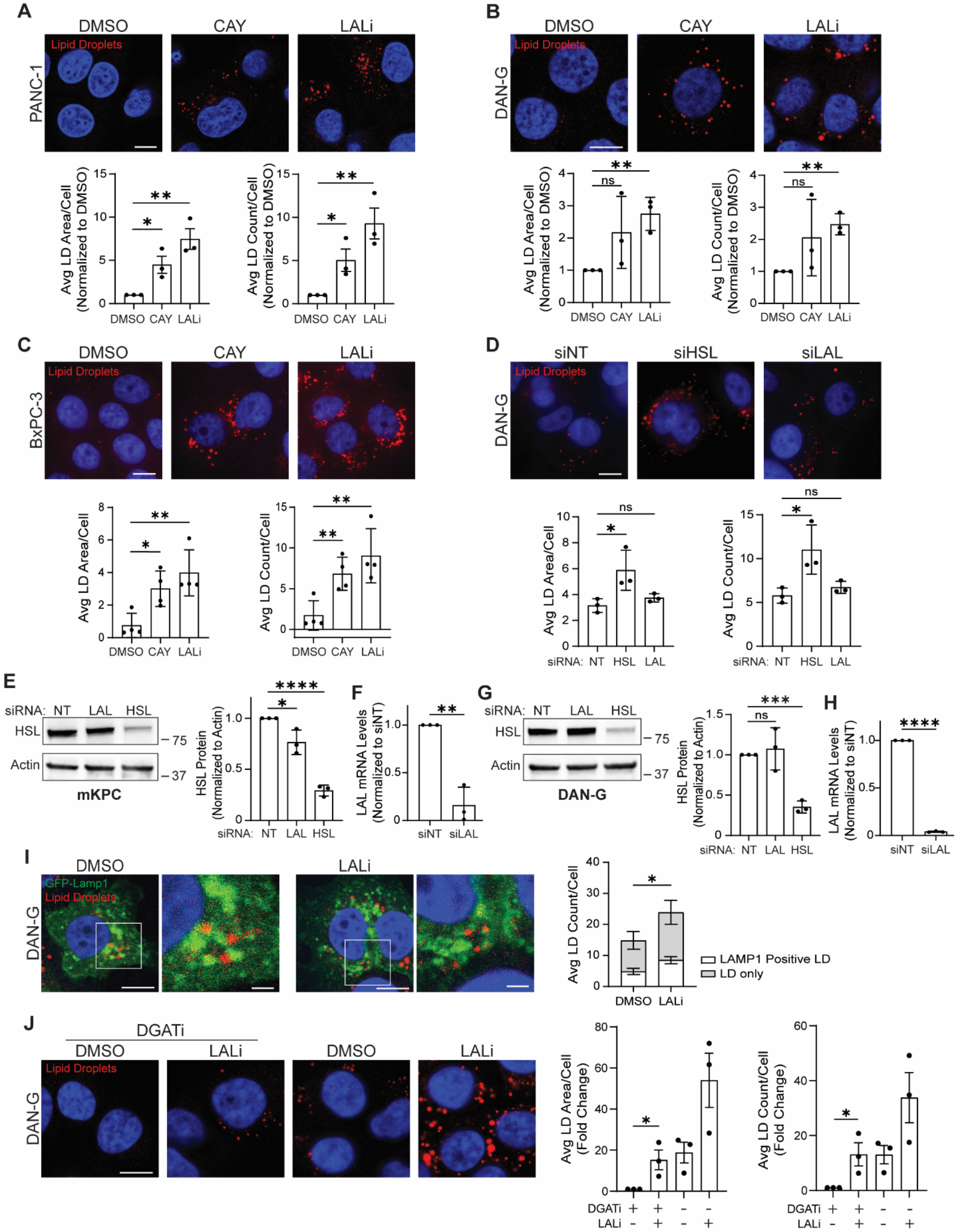

**Supplemental Figure S1. Lysosomal acid lipase promotes lipid droplet catabolism in pancreatic cancer cells.** Treatment of PANC-1 **(A)**, DAN-G **(B)**, or BxPC-3 **(C)** cells with DMSO (vehicle control), CAY10499 (CAY, 10μM), LAListat1 (LALi, 50 μM), or CAY + LALi for 24h and lipid droplets (Oil Red O,red). Graphs show quantitation of average lipid droplet (LD) area per cell and average LD count per cell. **D)** DAN-G cells transfected with siRNA (control non-targeting (NT), HSL, or LAL) and stained for lipid droplets (red) and nuclei (blue). Quantitation of 10 fields per condition in each biological replicate shows average lipid droplet area or count per cell (A-D). **E-H)** Knockdown efficiency was determined by western blotting for HSL and quantitative RT-qPCR for LAL. Representative immunoblots and quantitation of three independent replicates to demonstrate the degree of HSL knockdown (HSL normalized to Actin loading control) in mKPC cells **(E)** and DAN-G cells **(G)**. LAL mRNA levels were measured by RT-qPCR from three independent replicates to confirm LAL knockdowns for mKPC cells **(F)** and DAN-G cells **(H)**. **I)** DAN-G cells transfected with LAMP1-GFP (lysosomes, green), treated with DMSO or 50μM LALi for 24h, and stained for lipid droplets (Oil Red O, red) and nuclei (blue). Quantitation of average number of lipid droplets within LAMP1 positive lysosomes per cell of three independent biological replicates (n=10 fields per condition for each experiment). Statistical significance was calculated for LAMP1-positive lipid droplets. **J)** DAN-G cells treated with DGATi 1 and 2 at the same time as DMSO or 50μM LALi for 24h and stained for lipid droplets (red) and nuclei (blue). Graphs indicate the mean +/- SD of 3-4 independent biological replicates. Statistical significance was determined by unpaired Student's t test. *$p<0.05$, **$p<0.01$, ***$p<0.001$, ****$p<0.0001$, ns = not significant. Scale bar = 10μm for all images except the zoom in fields (I) scale bar = 1μm.

Supplemental Figure S2

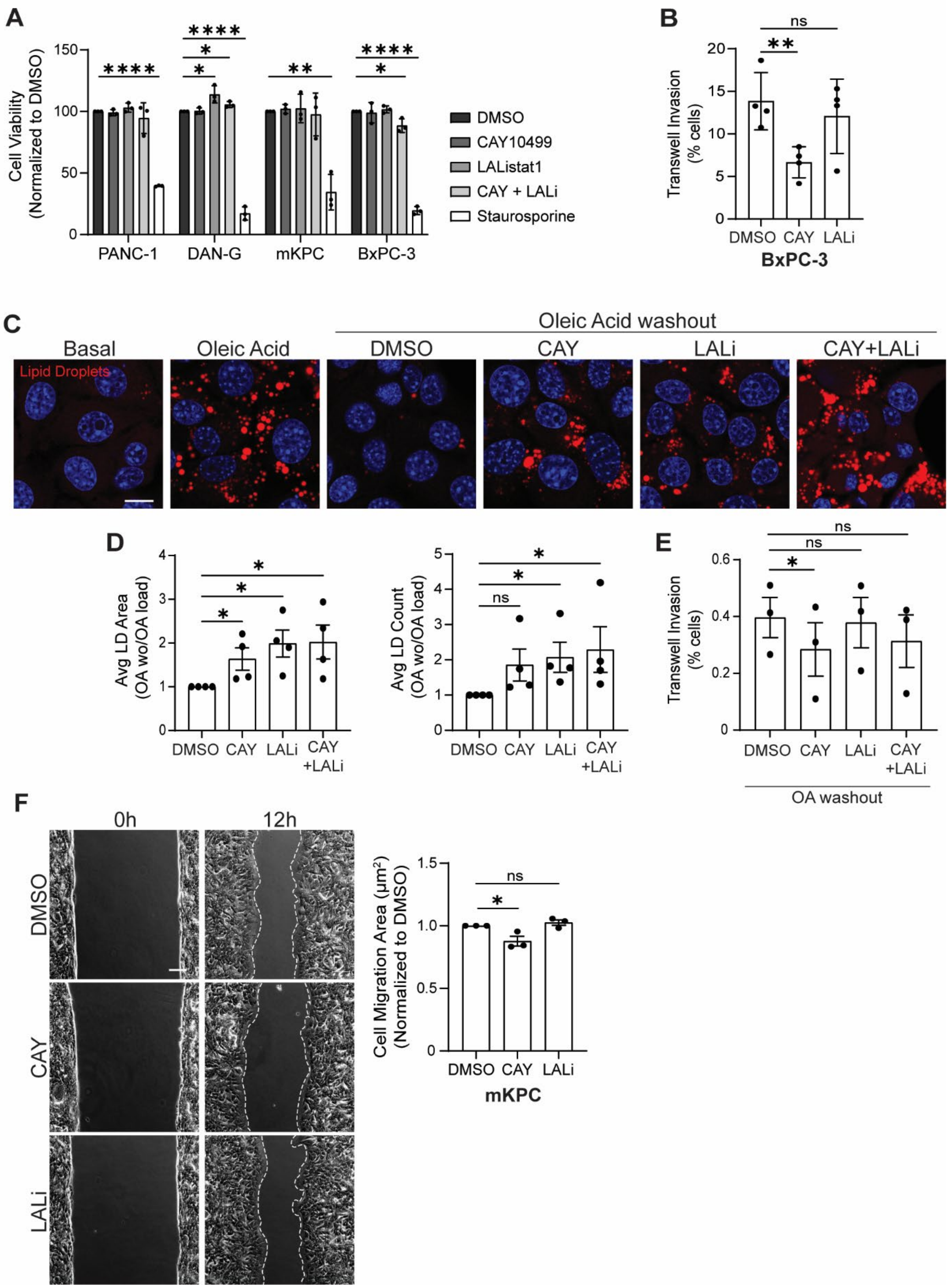

**Supplemental Figure S2. Lysosomal acid lipase does not regulate viability, 2D migration, or invasion in the context of excess oleic acid. A)** Lipase activity does not control cell viability in culture as measured by Cell Counting Kit-8 (CCK8) viability assay. Cells were treated with either DMSO, 10μM CAY, 50μM LALi, CAY+LALi, or Staurosporine (positive control, 5μM) for 24h prior to CCK8 measurement. **B)** Quantitation of BxPC-3 transwell cell invasion. Cells treated with DMSO, 10μM CAY, or 50μM LALi for 24h before replating in transwell chambers in the presence of inhibitors for an additional 24h. **C)** mKPC cells treated with oleic acid (OA, 200μM) for 24h, washed out and fresh media added with either DMSO, 10μM CAY, 50μM LALi, or CAY+LALi for 24h. Representative images, stained for lipid droplets (Oil Red O, red) and nuclei (blue). **D)** Quantitation of average lipid droplet area per cell and average number of lipid droplets per cell, all normalized to OA load condition. Three independent biological replicates, n=10 fields imaged per experiment for each replicate. **E)** Quantitation of transwell cell invasion assay for mKPC cells treated with oleic acid (OA, 200μM for 24h), washed out and fresh media added with either DMSO, 10μM CAY, 50μM LALi, or CAY+LALi for 24h. p values were calculated using paired Student's t test. **F)** Representative images of wound healing cell migration at 0h and 12h in mKPC cells treated with DMSO, 10μM CAY, or 50μM LALi for 24h prior to wound healing. Quantitation of cell migration area ($\mu m^2$) (n=3 fields per timepoint per condition for each experiment). Graphs indicate mean +/- SD for 3-4 independent biological replicates and statistical significance was determined by unpaired Student's t test. *$p<0.05$, **$p<0.01$, ****$p<0.0001$, ns = not significant. Scale bar = 10μm.

Supplemental Figure S3

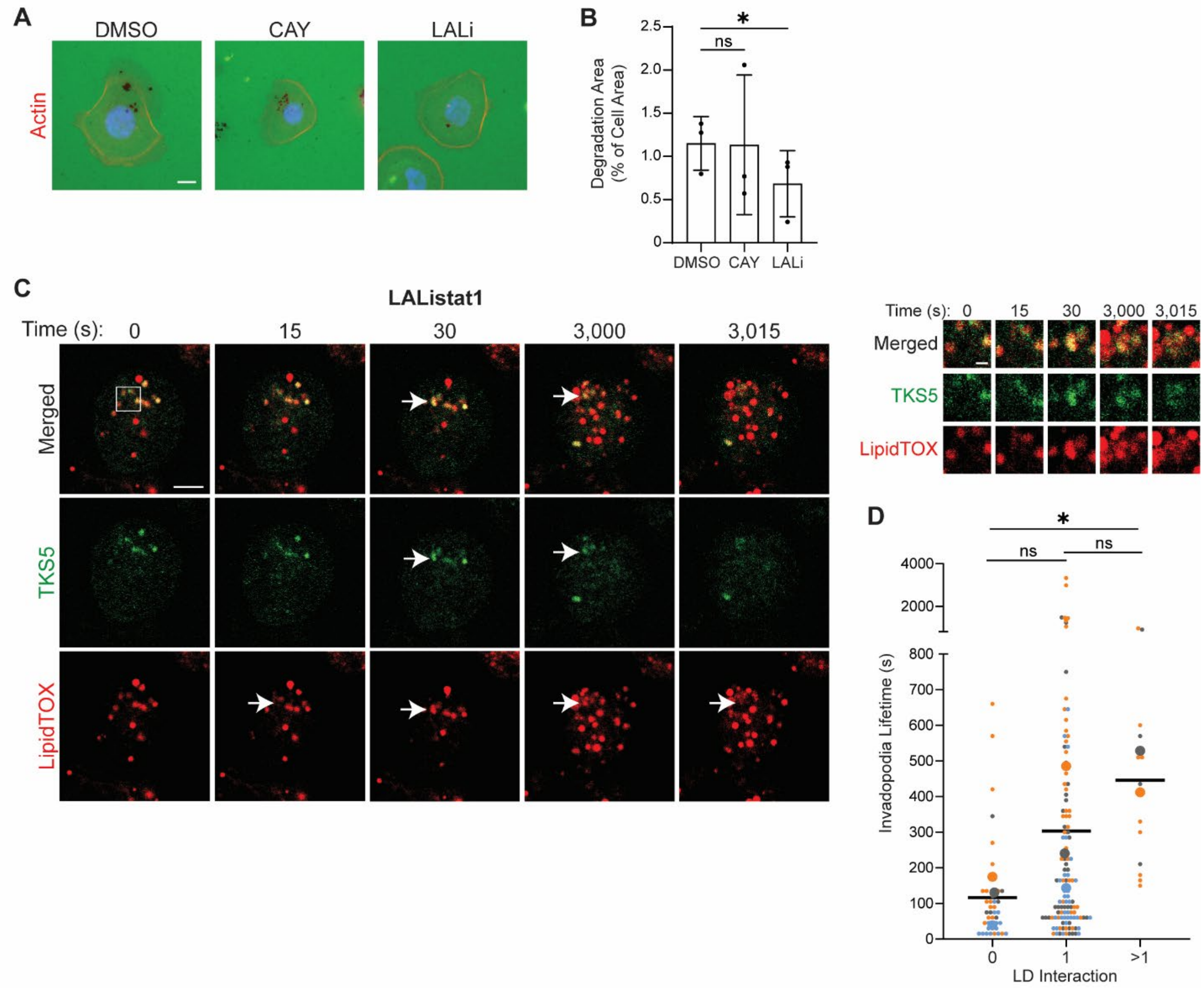


**Supplemental Figure S3. Lysosomal acid lipase promotes invadopodia-mediated extracellular matrix degradation. A)** BxPC-3 cells were pre-treated with either DMSO, 10μM CAY, or 50μM LALi for 24h. Cells were replated on Oregon green-conjugated gelatin coated coverslips for another 24h incubation with the same inhibitor treatments before fixing and staining with Hoechst (nuclei, blue) and phalloidin (Actin, red). Scale bar = 10μm. **B)** Quantitation shows average degradation area per cell, normalized to cell area. 10 fields were imaged and quantified per experiment in each of three independent biological replicates. Graphs indicate the mean +/- SD. **C)** DAN-G cells transfected with TKS5-GFP (green), treated with 50μM LALi for 24h, plated on unlabeled gelatin coated coverslips for 5h and stained with LipidTOX Deep Red (red) to visualize invadopodia (TKS5) and LD colocalization. Live cell imaging was performed, and representative images are shown from the indicated time points. Arrows point to TKS5 and lipid droplet colocalization. Invadopodia and lipid droplets were maintained in frames not shown between 30 and 3,000s. Magnified images show lipid droplet interaction with invadopodia over time. Scale bar = 5μm; magnified image scale bar = 1μm. **D)** TKS5-GFP positive invadopodia lifetime was quantitated and graphed based on number of lipid droplet interactions. Data points indicate all individual invadopodia imaged over three independent experiments. Statistical significance was determined by unpaired Student's t test **(B)** and paired Student's t test **(D).** *$p<0.05$, ns = not significant.

Supplemental Figure S4

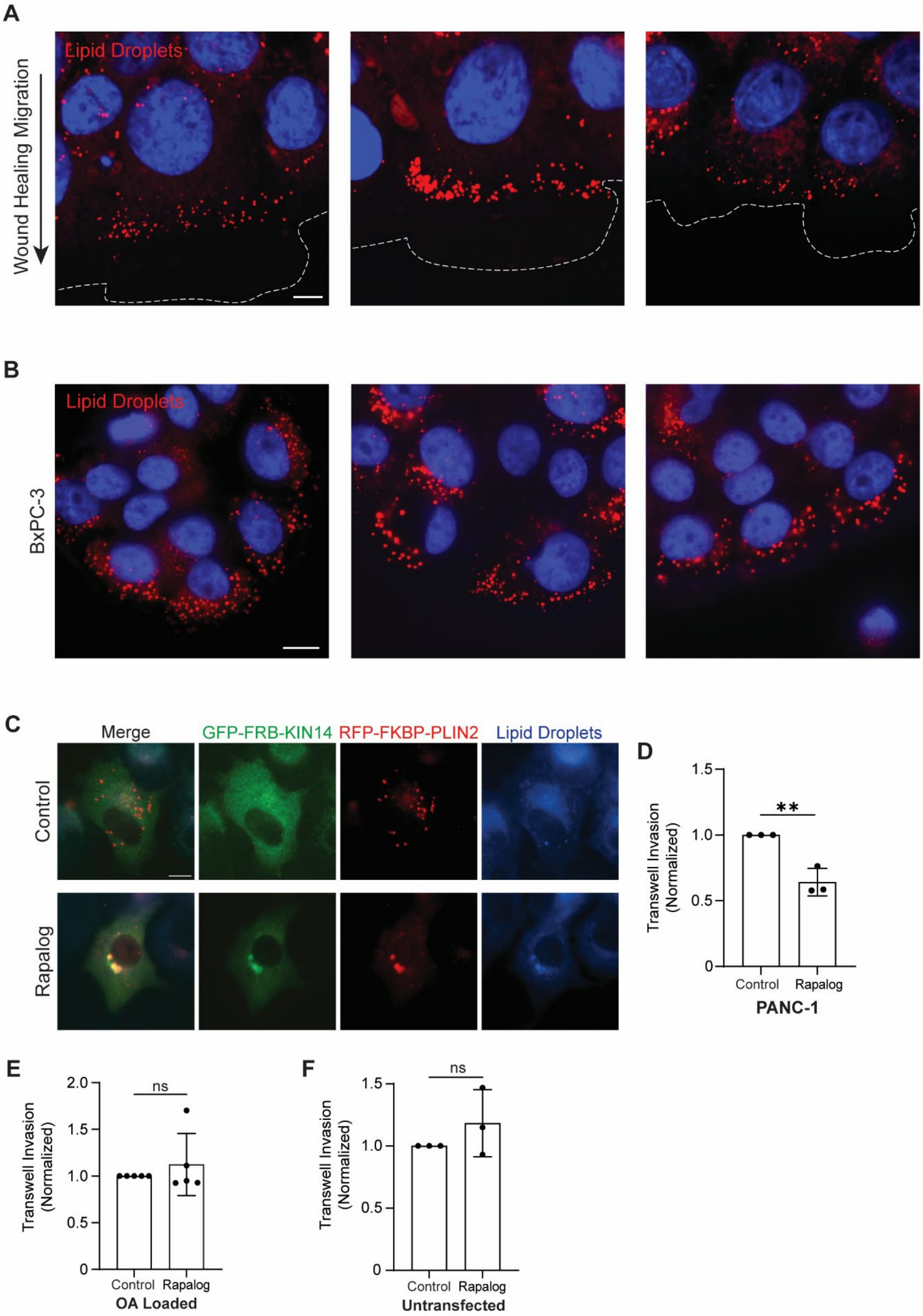

**Supplemental Figure S4. Lipid droplet localization is critical for tumor cell invasion. A)** BxPC-3 cells transfected with BODIPY (pseudo-colored red) to visualize lipid droplets localized to the cell periphery during cell migration. Scale bar = 5μm. **B)** BxPC-3 cells stained with ORO to visualize lipid droplets at the cell periphery. Scale bar = 10μm. **C)** PANC-1 cells transfected with RFP-FKBP-PLIN2 and GFP-FRB-KIN14 and treated with or without Rapalog A/C Heterodimerizer (10nM) for 4h. Lipid droplets were stained with MDH (blue) to show lipid droplet perinuclear clustering after Rapalog treatment. **D)** PANC-1 cells transfected and treated as in (A) were seeded in a transwell assay for 48h. Transfected cells on the top (non-invaded) and bottom (invaded) of the filter were scored to calculate percent invasion. **E)** mKPC cells were transfected and treated with 10nM Rapalog after oleic acid (OA, 200μM) loading and washout and indicated no change in invasion after 24h. Only transfected cells indicating lipid droplet trafficking were scored to calculate the percentage of cells invaded (B-C). **F)** Untransfected mKPC cells were treated with and without 10nM Rapalog to show that the Rapalog alone does not change transwell cell invasion. n=10 fields per condition. Graphed data represent the mean ± SD for at least three independent biological replicates. p values were calculated using unpaired Student's t test. $^{**}p < 0.01$, ns = not significant.

Supplemental Figure S5

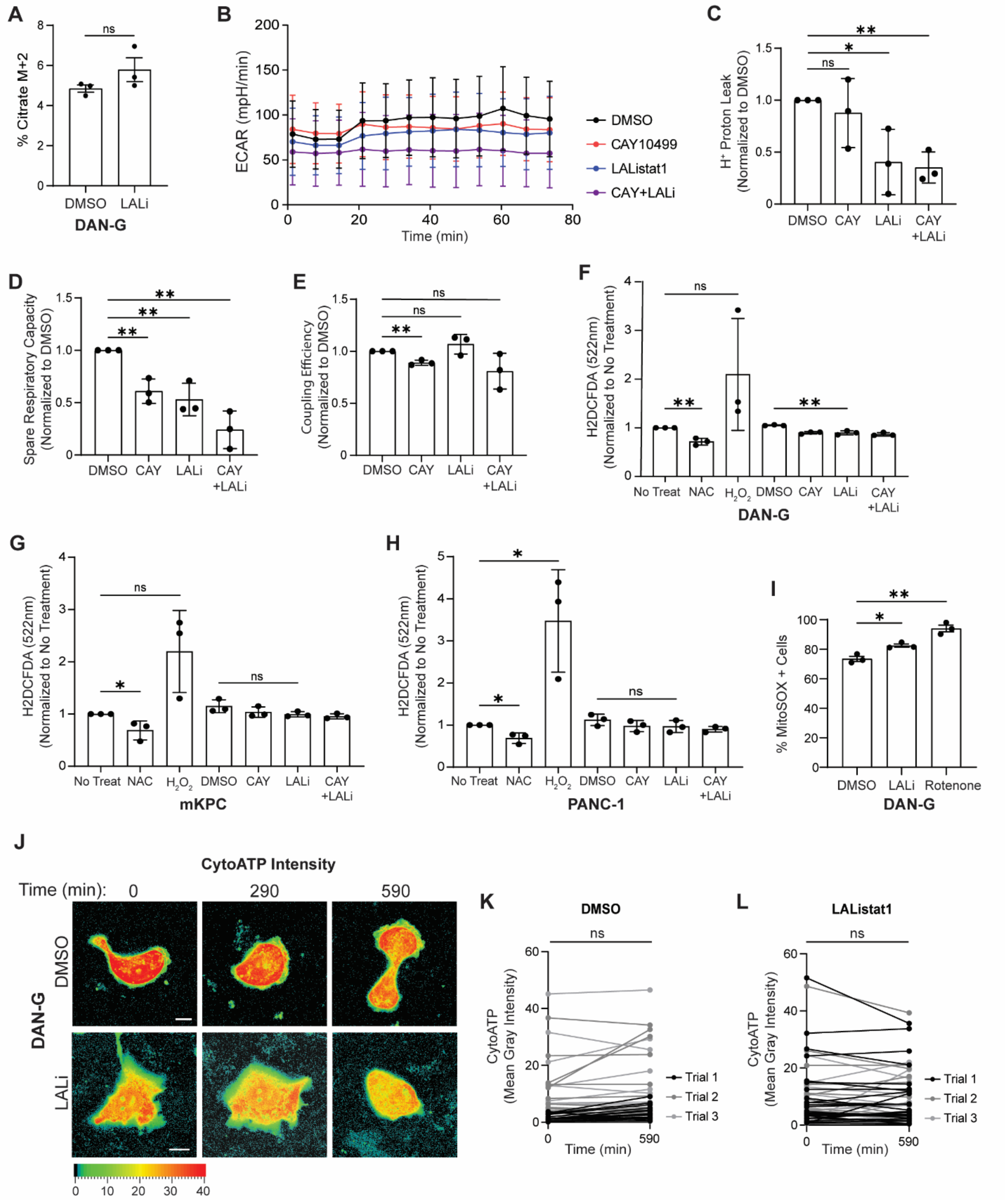


**Supplemental Figure S5. Cytosolic and lysosomal acid lipase activity regulate oxidative metabolism and ATP production. A)** Quantitation of palmitic acid tracing in

DAN-G cells treated with DMSO or 50μM LALi and measurement of percent citrate levels (M+2). **B-E)** Quantitation of average normalized ECAR from Seahorse analysis output for DAN-G cells treated with either DMSO, 10μM CAY, 50μM LALi, or CAY+LALi for 24h. Graphs represent data from three independent biological replicates with four wells averaged per condition. Data extracted from Figure 5A includes H+ Proton leak **(C),** Spare Respiratory Capacity **(D)** and Coupling Efficiency **(E)**. **F-H)** Reactive oxygen species (ROS) measurements using H2DCFDA for DCF fluorescence following excitation 493nm and emission of 522nm DAN-G **(F),** mKPC **(G)** and PANC-1 **(H)** cells were treated with DMSO, 10μM CAY, 50μM LALi, or CAY+LALi for 24h, N-acetylcysteine (5mM, negative control) or $H_2O_2$ (1mM, positive control) for 15min prior to H2DCFDA addition and measurement. **I)** Mitochondrial ROS measurements using MitoSOX by flow cytometry analysis on DAN-G cells treated with DMSO, 50μM LALi for 24h, or 1μM Rotenone (positive control) for 2h. For A-I, graphed data represents the mean ± SD for at least three independent biological replicates. **J)** DAN-G cells transfected with cyto-iATPSnFR1.0. At t=0, DMSO or 50μM LALi were spiked in and cells were imaged over 590min. Fluorescence intensity of cyto-iATPSnFR1.0 (CytoATP) is pseudo-colored via Look-Up table (LUT) heatmap, with red indicating relatively higher ATP levels. Scale bar = 10μm **K-L)** Quantitation from live cell imaging in (J) of cyto-iATPSnFR1.0 mean gray intensity at TKS5 positive puncta per cell at time 0 and 590min treated with DMSO **(K)** or LALi **(L)**. Each line represents the average invadopodia fluorescence intensity of one cell. Statistical significance was determined by unpaired Student's t test. *p<0.05, **p<0.01, ns = not significant.

Supplemental Figure S6

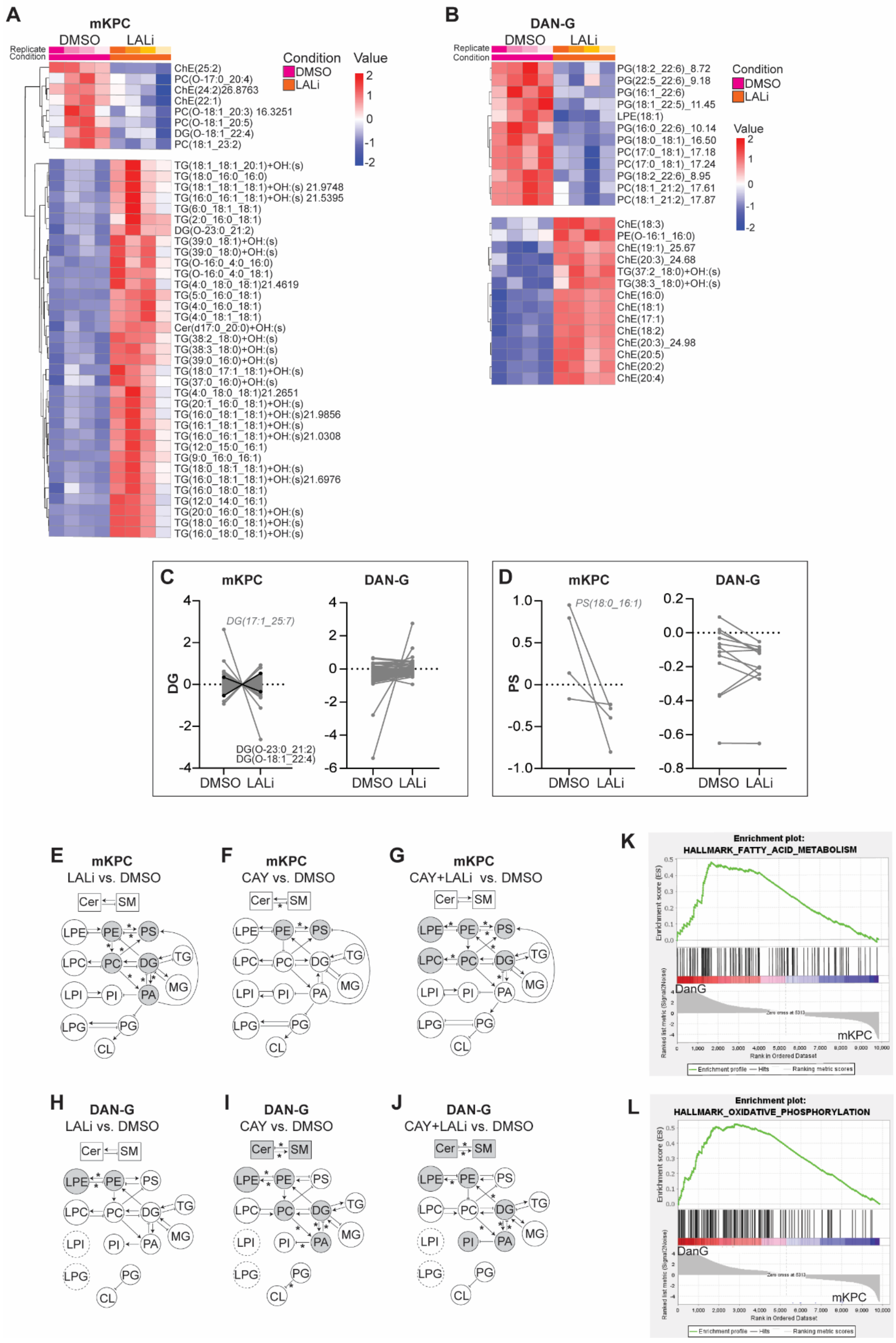

**Supplemental Figure S6. Lysosomal lipolysis modulates membrane lipid composition and tension.** DAN-G and mKPC cells were treated with either DMSO, 10μM CAY, 50μM LALi, or CAY+LALi for 24h, pelleted, and analyzed by lipidomics. Heatmap of specific species from lipidomics output for mKPC ($p \leq 0.05$) cells **(A)** and DAN-G ($p \leq 0.1$) cells **(B)** treated with DMSO or 50μM LALi generated through RStudio for four independent biological replicates. **C-D)** Specific paired lipid species comparison from mKPC and DAN-G cells treated with DMSO or 50μM LALi of diglycerides (DG) **(C)** and phosphatidylserine (PS) **(D)**. Black lines indicate species with statistically significant changes. **E-J)** Diagrams of BioPAN pathway analysis for mKPC **(E-G)** and DAN-G **(H-J)** cells treated with either 50μM LALi **(E,H),** 10μM CAY **(F,I)**, or CAY+LALi **(G,J)** compared to DMSO treatment. Arrows represent significant ($p \leq 0.05$) interactions between lipid subclasses. Asterisks (*) represent significant $|Z|$ scores above 1.645. Gray shading represents active or suppressed status in BioPAN rendering from lipids processed output after separate conversion through LipidLynxX. **K-L)** Enrichment plots for DAN-G vs. mKPC cells of MSigDb Hallmark gene set collections for Hallmark fatty acid metabolism (NES= 1.42, Nominal p-value= 0.0, FDR q-value= 0.47) **(K)** and Hallmark oxidative phosphorylation (NES=1.43, Nominal p-value= 0.0, FDR q-value= 0.49) **(L)**.

### Supplemental Movies

**Movie S1.** DAN-G cells expressing TKS5-GFP to label invadopodia and incubated with LipidTOX Deep Red to label lipid droplets. Lipid droplets appear prior to the formation of invadopodia and persist after invadopodia disassembly.

**Movie S2.** DAN-G cells, treated with LAListat1, expressing TKS5-GFP to label invadopodia and incubated with LipidTOX Deep Red to label lipid droplets.

**Movie S3.** mKPC cells under basal conditions incubated with LipidTOX Deep Red to label lipid droplets and Lysotracker Green to label lysosomes. Lipid droplets are present at the cell periphery of the actively migrating mouse pancreatic cancer cells.

**Movie S4.** mKPC cells under basal conditions, treated with LAListat1, incubated with LipidTOX Deep Red to label lipid droplets and Lysotracker Green to label lysosomes. Lipid droplets are present the cell periphery of the actively migrating mouse pancreatic cancer cells.

**Movie S5.** PANC-1 cells after oleic acid loading and washout, incubated with LipidTOX Deep Red to label lipid droplets, Lysotracker Blue to label lysosomes, and MitoTracker Green to label mitochondria. Lipid droplets localize to and are quickly catabolized at the cell periphery of the actively migrating pancreatic cancer cells.

**Movie S6.** PANC-1 cells after oleic acid loading and washout with LAListat1 treatment, incubated with LipidTOX Deep Red to label lipid droplets, Lysotracker Blue to label lysosomes, and MitoTracker Green to label mitochondria. Lipid droplets are present at the cell periphery of the actively migrating pancreatic cancer cells and accumulate in the presence of LAListat1.

**Movie S7-9.** DAN-G cells transfected with TKS5-mCherry to label invadopodia and cyto-iATPSnFR1.0 to label ATP, with DMSO spiked in right before start of the movie. ATP levels at invadopodia do not change in the presence of DMSO treatment.

**Movie S10-12.** DAN-G cells transfected with TKS5-mCherry to label invadopodia and cyto-iATPSnFR1.0 to label ATP, with LAListat1 spiked in right before start of the movie. ATP levels at invadopodia decrease in the presence of LAListat1 treatment.
